# P3 site-directed mutagenesis: An efficient method based on primer pairs with 3’-overhangs

**DOI:** 10.1101/2024.10.18.615663

**Authors:** Negar Mousavi, Ethan Zhou, Arezousadat Razavi, Elham Ebrahimi, Paulina Varela-Castillo, Xiang-Jiao Yang

## Abstract

Site-directed mutagenesis is a fundamental tool indispensable for protein and plasmid engineering. An important technological question is how to achieve the efficiency at the ideal level of 100%. Based on complementary primer pairs, the QuickChange method has been widely used, but it requires significant improvements due to its low efficiency and frequent unwanted mutations. An alternative and innovative strategy is to utilize primer pairs with 3-prime overhangs, but this approach has not been fully developed. As the first step towards reaching the efficiency of 100%, we have optimized this approach systematically (such as use of newly designed short primers, test of different Pfu DNA polymerases and modification of PCR parameters) and evaluated the resulting method extensively with over 100 mutations on 12 mammalian expression vectors, ranging from 7.0-13.4 kb in size and encoding ten epigenetic regulators with links to cancer and neurodevelopmental disorders. We have also tested the new method with two expression vectors for the SARS-COV-2 spike protein. Compared to the QuickChange method, the success rate has increased substantially, with an average efficiency of ∼50%, with some at or close to 100 percent, and requiring much less time for engineering various mutations. Therefore, we have developed a new site-directed mutagenesis method for efficient and economical generation of various mutations. Notably, the method failed with a human KAT2B expression plasmid that possesses extremely GC-rich sequences. Thus, this study also sheds light on how to improve the method for developing ideal mutagenesis methods with the efficiency at ∼100% for a wide spectrum of plasmids.

## Introduction

Whole-genome and exome sequencing have yielded libraries of germline or somatic mutations in various diseases, such as genetic disease-linked mutations listed in the recently established Clinvar database [1] and cancer-associated mutations documented in the cBioportal and COSMIC databases [2,3]. Many of these mutations are missense and their functional impact is often difficult to predict, so an important question is to ascertain whether these mutations are causal. To establish the pathogenicity, one standard approach is to engineer the mutations for functional analysis with molecular and cell-based assays *in vitro,* followed by generation and analysis of model organisms with equivalent mutations *in vivo*. For the assays *in vitro*, site-directed mutagenesis of the affected genes is often employed. Furthermore, site-directed mutagenesis is frequently employed to generate mutants for modelling evolution of viruses (such as SARS-COV-2) and assessing the importance and impact of new mutations acquired during evolution. Thus, site-directed mutagenesis is a basic tool for biomedical research.

Since the initial report of this invention by the Nobel Laureate Michael Smith and his colleagues in 1978 [4], the technique has been widely used for engineering gene mutations at specific sites of phage or plasmid DNA *in vitro*. For introducing site-specific mutations into plasmid DNA, various protocols were developed to enhance the efficiency, including insertion of an F1 replication origin into a plasmid vector for isolation of uracil-containing single-stranded phagmid DNA as the mutagenesis template [5-8]. But the isolation of single-stranded phagmid DNA is an extra step, which is lengthy, not straightforward or even unsuccessful for some phagmid vectors. Overcoming such limitations, the QuickChange^™^ (or QuikChange^™^ as marketed by Agilent) site-directed mutagenesis method was developed to utilize double-stranded plasmids directly as DNA templates [9-11]. It is based on Pfu DNA polymerase-mediated PCR with a pair of complementary primers containing a given mutation (Fig. 1A). This is followed by DpnI digestion, which selectively degrades the wild-type parental plasmid isolated from a Dam-positive *E. coli*, where methylation at GATC sites occurs for recognition by DpnI. Phosphorylation of primers was initially required for a ligation step [9-11], but this requirement was eliminated in the QuikChange® protocol.

**Fig. 1.**
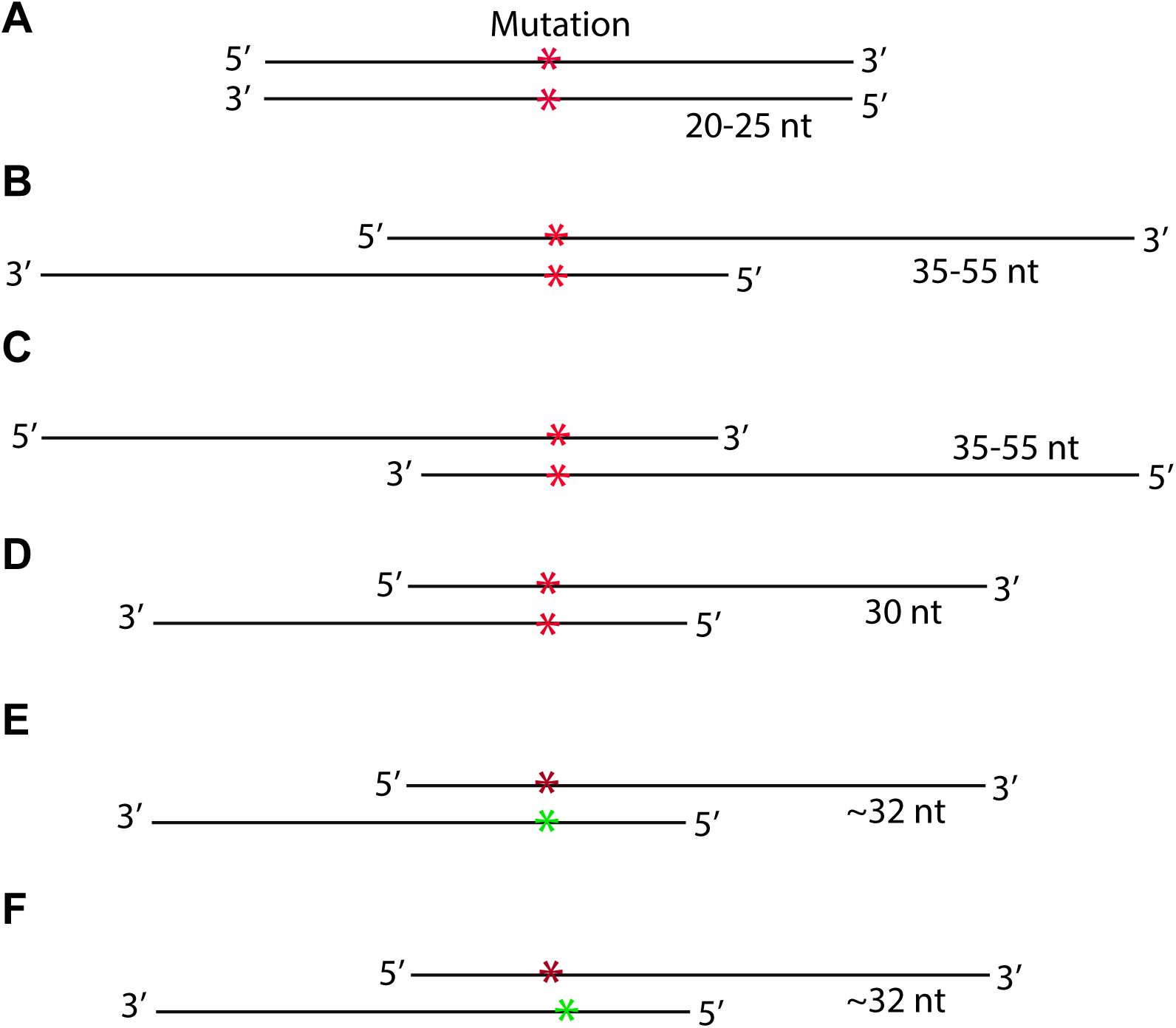
Primer design for different site-directed mutagenesis methods. A. Completely complementary primer pair for the Quickchange^TM^ mutagenesis method [9,10]. Two red asterisks mark the mutation sites within the primer pair. The primers are 20-25 nucleotides (nt) in length and completely complementary to each other. B. Partially complementary primer pair with 3’-overhangs. Unlike those in panel A, such primers can anneal to newly synthesized strands and initiate synthesis of the complementary strand until reaching the 5’end of the opposite primer (see Fig S1A-B). Liu & Naismith designed primers with the average about 45 nucleotides (ranging from 35 to 55 nucleotides) in length, but special care was recommended to select the primers [13]. C. Partially complementary primer pair with 5’-overhangs. Such primers can anneal to newly synthesized strands serve as templates for very limited synthesis of the complementary strand until reaching the 5’end of the opposite primer. Another drawback is that the DNA polymerase can initiate 5’→3’ synthesis on the primer pair and make it similar to that in panel A. Thus, this design suffers similar problems as the completely complementary primer pair (A). D. Shorter primer pair with 3’-overhangs. We have simplified the primer design and reduced the length of the two primers to 30 nucleotides, with 9-bp complementary regions flanking a single-nucleotide mutation site. No other special care is required for selection of the length of the primers. Compared to the design in panel B, the shorter primers reduce cost and minimizes unwanted mutations introduced by primer impurity resulting from chemical synthesis. Notably, this is a major source of unwanted mutations when high-fidelity DNA polymerases are used for PCR-based site-directed mutagenesis. E-F. The two primers for engineering the mutation can carry a mismatch at the same position as indicated with red and green asterisks in panel D. Moreover, the primers can even carry different mutations at nearby positions, as illustrated with red and green asterisks in panel E. In these cases, the primers are slightly longer than 30 nucleotides, with ∼10-bp complementary regions encompassing the mismatched mutation sites. Thus, in such cases, two different mutants can be generated with a single pair of primers. G. Two primers carry different mutations at their 3’-arms, as illustrated with red and green asterisks. A slightly different version is to have one mismatch in the complementary region and a second mutation in one of the 3’-arms. This also illustrates an additional advantage that primer pairs with 3’-overhangs possess when compared to the completely complementary primer pairs (A).

This simplified protocol has been widely used, but difficulties have been encountered, requiring solutions case by case [12,13]. In agreement with this, the method has demonstrated reasonable efficacy for generating mutants of various histone-modifying enzymes in our laboratory [14-17], but it took considerable efforts to generate various mutants, often requiring multiple trials and different optimization efforts. There are three underlying limitations with the protocol: 1) the complementary primers self-anneal and generate primer-primer dimers during PCR, thereby decreasing the amplification efficiency; 2) newly synthesized plasmid DNA is ‘nicked’ and thus unsuitable as a template for further amplification; and 3) unwanted mutations at the primer sites, due to impurity of primers and strand-displacement by Pfu.

As illustrated in Fig. 1B, an alternative approach is to utilize a pair of partially complementary primers with two 3’-protruding ends, thereby overcoming limitations 1) and 2) mentioned above. In comparison, primers with 5’-ovehangs (Fig. 1C) cannot overcome such limitations. The simple but innovative approach (Fig. 1B) was initially implemented by Zheng, Baumann & Reymond [12]. One drawback of this initial study was that the 5’arm is only 3-5 nucleotides long for some primers, which leads insufficient annealing and priming from newly synthesized strands. As a result, this only partially address limitation 1) mentioned above (Fig. 1B) [12]. Another drawback is that only restriction digestion at coupled secondary sites, instead of DNA Sequencing, was used to assess the mutagenesis efficiency, thereby leaving the uncertainty about the real sequences at the mutation sites of the selected clones [12]. This method was later refined by Liu & Naismith, leading to its broader application and recognition [13]. Since the method was developed and only tested for 5-6 kb plasmids [13], an important question is how it performs with larger plasmids and those with more complex sequences, as site-directed mutagenesis becomes more challenging with larger plasmids [18-20].

To develop this innovative method further, we have optimized it systematically in the following six aspects, which together represent significant advancement and make the resulting method efficient, versatile and economical. First, we used short primers only ∼30-nucleotides long (Fig. 1D). Such shorter primers are expected to reduce unwanted mutations at the primer sites, as longer primers increase the likelihood of errors during synthesis. Since the coupling efficiency of each nucleotide addition is slightly below 100%, longer oligonucleotides accumulate, such as those used in the previous study [13], more mutations with each synthesis cycle. This not only reduced the costs but also minimized potential unwanted mutations introduced by intrinsic primer impurities from chemical synthesis. This is because oligonucleotide synthesis, each cycle of nucleotide addition has an efficiency slightly below 100%. Thus, from chemical synthesis, shorter primers are typically of higher quality than longer ones and reduce unwanted mutations caused by some incorrect primer molecules existing in unpurified primers as impurities. Second, instead of one mutation per primer pair, two different mutations were introduced with a single pair of primers when necessary (Fig. 1E-F). These two mutations can even be on the 3’-arms of the primers (Fig. 1G). Third, as it displays ∼6 folds higher fidelity than Taq DNA polymerase [21,22], Pfu DNA polymerase was used in the previous two studies [12,13]. We have utilized an improved version of Pfu, known as PfuUltra, composed of Pfu DNA polymerase and a dUTPase to improve PCR performance [23]. Compared to Pfu itself, the error rate was reduced further by ∼3 folds. Fourth, to speed up PCR amplification, we tested two fast versions of Pfu DNA polymerase, PfuUltra II and Pfu-fly. The former is 70-80% faster than PfuUltra, while Pfu-fly was advertised to be at least 5 times faster than Pfu itself and display much higher fidelity than PfuUltra (∼108 folds higher than Taq polymerase). Fifth, we used a fast PCR program, a much small reaction volume of 5-10 μl (instead of 50 μl) and a fresh PCR tube for DpnI digestion. For this, DpnI was added to the reaction mixture after PCR and the mixture was then transferred to a fresh PCR tube for incubation at 37°C. This step helps eliminate contamination of undigested template plasmid molecules adhered to the inner wall of the old PCR tube. This is especially important when the colony number is low after transformation. Finally, we adopted an efficient and economical in-house protocol to prepare competent cells for transforming 2-5 μl of a DpnI-digested reaction mixture. Notably, these competent *E. coli* cells can be prepared in large batches and stored in liquid nitrogen tanks for up to 12 years, with minimal loss of competence. To distinguish it from the QuickChange^™^ protocol, we refer to this systematically optimized method as P3 (<u>p</u>rimers with <u>3</u>’-protruding ends) site-directed mutagenesis.

To assess its versatility, we have evaluated this new method with >100 mutations on a dozen of expression vectors, ranging in size from 7.0-13.4 kb, all of which are notably larger than the 5-6 kb size tested in the previous study [13]. Compared to the QuickChange^™^ method, the success rate has increased significantly, reaching an average efficiency of >50%, at or close to 100% in some cases, and requiring less time for engineering various mutations. Typically, only 2-3 colonies need to be analyzed per mutagenesis reaction. As a result, a skillful trainee can generate a few dozens of mutants within a week. Moreover, Pfu-fly outperforms PfuUltra, in terms of PCR length, colony yield and overall success, although it frequently causes unwanted deletions and insertions at the primer or mutation sites. The reagent cost per mutagenesis reaction is reduced to ∼$0.5-1, excluding costs for primers and plasmid sequencing. The strategies shown in Fig. 1E-G reduce primer and sequencing costs further by ∼50%. Intriguingly, among all plasmids that we tested, the one for BRPF3 was significantly more challenging than the others. Thus, in this study, we have developed an optimized mutagenesis method and enhanced its efficiency, versatility and cost-effectiveness. The extensive experience with different vectors should facilitate rapid and efficient mutagenesis of other plasmids. This study also sheds light on further improvements to develop ideal site-directed mutagenesis methods with the efficiency at or close to 100% for a broad variety of plasmids. Thus, this work nicely extends and complements the previous two studies [12,13].

## Results

### Optimizing P3 site-directed mutagenesis for generating BRPF1 clinical variants

BRPF1 (bromodomain- and PHD finger-containing protein 1) was initially identified and cloned as BR140 (bromodomain protein with an estimated molecular weight of 140 kDa) [24]. As illustrated in Fig. 2A, it possesses multiple domains for epigenetic regulation, including the N-terminal part for interacting specifically with lysine acetyltransferase 6A (KAT6A) and its paralog KAT6B [25]. The interaction is required for activation of these two acetyltransferases. In addition, BRPF1 possesses two EPC (Enhancer of Polycomb)-like modules, the second of which mediates interaction with ING4 (or its paralog ING5) and MEAF6. Thus, BRPF1 is a scaffold protein important for forming and activating multiple tetrameric acetyltransferase complexes composed of KAT6A (or KAT6B), ING4 (or ING5) and MEAF6 [25]. Moreover, BRPF1 possesses three different domains for recognition of nucleosomes, including a PZP module, a bromodomain and a PWWP domain (Fig. 2A) [25]. The PZP module is composed of two PHD fingers flanking a zinc knuckle, with the first PHD finger for recognition of the unmodified N-terminus of nucleosomal histone H3 and the second PHD finger function for binding to the DNA backbone of the nucleosome. In line with these multiple domains important for epigenetic regulation, recent studies have linked *BRPF1* mutations to a novel neurodevelopmental disorder [15,16,26-28]. Moreover, according to the Clinvar database [1], additional mutations have been identified in patients. While some of the identified *BRPF1* mutations have been molecularly characterized [15,16,26], many more remain to be analyzed to establish their pathogenicity.

**Fig. 2.**
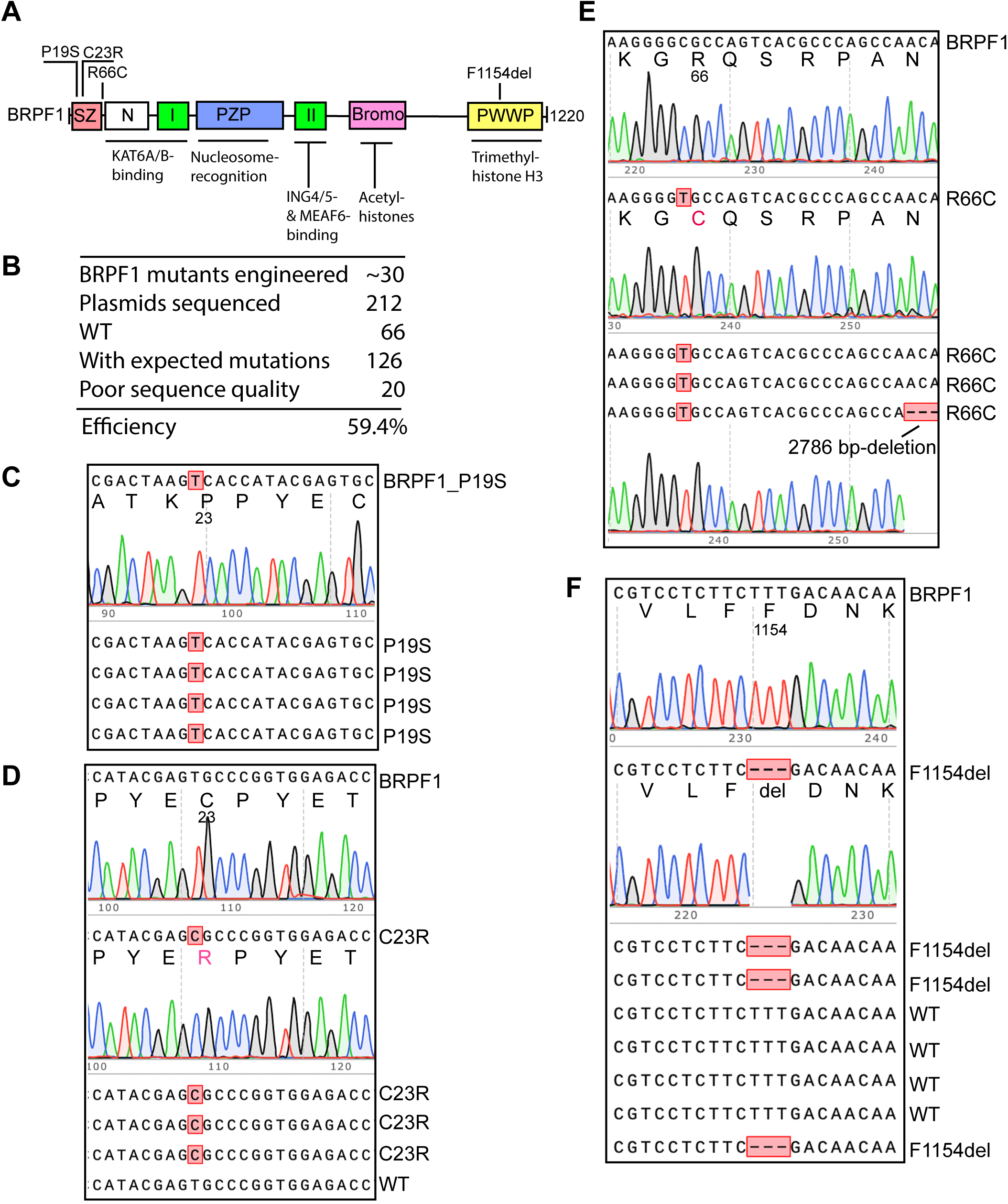
Efficiency of P3 site-directed mutagenesis to engineer BRPF1 mutants. A. Domain organization of BRPF1. It is composed of an Sfp1-specific zinc finger (SZ), an N-terminal domain (N) required for specific interaction with lysine acetyltransferase 6A (KAT6A) and its paralog KAT6B [25], two EPC (Enhancer of Polycomb)-like motifs (I and II), a PHD-zinc buckle-PHD (PZP) module, a bromodomain and a PWWP domain [25]. For simplicity, only four missense variants are illustrated, with two located to the SZ region (P19S and C23R), one between the SZ and the N domain (R66C) and one at the PWWP domain (F1154del). B. Efficiency of the mutagenesis method. Among 212 plasmids sequenced, 20 could not be sequenced, perhaps due to wrong plasmids (such as the lack of primer sequence), poor plasmid quality or Sanger sequencing issues. To be conservative, they were all counted as incorrect mutant candidates for efficiency calculation. Thus, the mutagenesis efficiency would be higher if the failed cases due to Sanger sequencing issues were eliminated from calculation. C. Sanger sequence analysis of 5 plasmids resulting from engineering the P19S mutant. All five were correct, so the mutagenesis efficiency was 100%. D. Sanger sequence analysis of 6 plasmids resulting from generating the C23R mutant. Four are correct and two are wild-type, so the mutagenesis efficiency was 4/6 (66.7%). E. Sanger sequence analysis of 7 plasmids resulting from engineering the R66C mutant. Two could not be sequenced (not shown here). Four contain the mutation but one of them possesses a ∼2.8-kb deletion, so the mutagenesis efficiency was 3/7 (42.9%). F. Sequence chromatograms of 9 plasmids analyzed for generating the F1154del mutant. Four are correct, four are wild-type and one could not be sequenced (not shown here), so the mutagenesis efficiency was 4/9 (44.4%).

In this regard, site-directed mutagenesis serves as a fundamental tool to engineer the mutants for functional analysis in biochemical and cell-based assays. Previously, we utilized the QuickChange^™^ method (Fig. 1A) to generate BRPF1 mutants for molecular characterization [15,16]. While the technique demonstrated considerable efficacy in experiments carried out by various trainees in the laboratory [14-17], it took considerable efforts to generate the mutants (especially for new trainees), often requiring multiple trials. Such issues have also been reported by others [12,13]. Despite its wide use in different laboratories, the QuickChange^™^ method suffer from at least two constraints. First, the primer pairs are completely overlapping, so there is a propensity for self-annealing to form primer dimers during PCR. Second, as the newly synthesized DNA strands are ‘nicked,’ they are unsuitable for use as templates for further amplification. As such, we sought an alternative approach to generate BRPF1 mutants more efficiently. Related to this, we noticed the innovative primer design strategy described in two published reports [12,13]. The strategy employs primer pairs with 3’-protruding ends (Fig. 1B) and addresses the two aforementioned limitations intrinsic to the completely complementary primers used in the QuickChange™ method (Fig. 1A). Thus, we utilized this new primer-designing strategy (Fig. 1B) to generate BRPF1 mutants derived from the Clinvar database. Notably, the size of the expression plasmid for HA-tagged BRPF1 is 9.0 kb and much larger than the 5-6 kb plasmids tested in the previous report [13].

To adopt this innovative strategy (Fig. 1B), we systematically optimized the resulting mutagenesis method in the following five aspects initially. First, we tested different primer length and utilized primers as short as 30 nucleotides, which is significantly shorter than those reported by others (Fig. 1D) [12,13]. Second, instead of one mutation per primer pair, two different mutations were introduced with a single pair of primers when necessary (Fig. 1E-F). Third, we utilized PfuUltra, an improved version of Pfu compared to that used in the previous studies [12,13]. Fourth, we refined PCR parameters by reducing the amount of parental DNA to 10-15 ng and minimized the PCR reaction volume to 10 μl, instead of 50 μl as reported by others [12,13]. Moreover, after PCR amplification, DpnI digestion was carried out in a new PCR tube to avoid contamination from undigested plasmid molecules adhering to the inner wall of the previous PCR tube above the reaction mixture, which was especially important in cases when PCR amplification was inefficient and the resulting colony number was low. Finally, we adopted an in-house protocol to generate efficient competent cells to transform DpnI-digested PCR products. With this optimized method (i.e., P3 mutagenesis), we generated ∼40 BRPF1 mutant rapidly and economically, with an average efficiency close to 60% (Fig. 2B-F). Typically, per mutation, we only needed to send three plasmids from the resulting colonies for Sanger sequencing, so a trainee could generate a dozen of mutants easily within a week. By comparison, it took weeks (and sometimes even months) of trials and errors (for optimizing PCR conditions) to obtain a few mutants when the QuickChange™ method was used [15,16].

### P3 site-specific mutagenesis for engineering BRPF2 and BRPF3 mutants

Considering the success with engineering BRPF1 mutants, we extended the optimized method to generate mutants of two paralogs, BRPF2 (a.k.a. BRD1, Fig. 3A) and BRPF3 (Fig. 4A). At the amino acid sequence and domain organization levels, BRPF1 is highly homologous to BRPF2 and BRPF3. But KAT7 is a preferred partner of BRPF2 and BRPF3 [29-32], critical for acetylation of histone H3 at lysine 14 *in vitro* and *in vivo* [29,33]. By contrast, BRPF1 activates KAT6A and KAT6B to acylate histone H3 at lysine 23 [15,16]. Thus, BRPF1 is functionally different from its paralogs, BRPF2 and BRPF3. BRPF1 is linked to developmental disorders [15,16,26-28], so a logical question is whether BRPF2 and BRPF3 also play a role in developmental disorders. According to the Clinvar database [1], there are *BRD1* mutations linked to genetic diseases. These mutations have not been molecularly characterized, so it is important to engineer them and assess their pathogenicity. As the first step, we have engineered some of these mutants (Fig. 3), with the average mutagenesis efficiency of 54% (Fig. 3B).

**Fig. 3.**
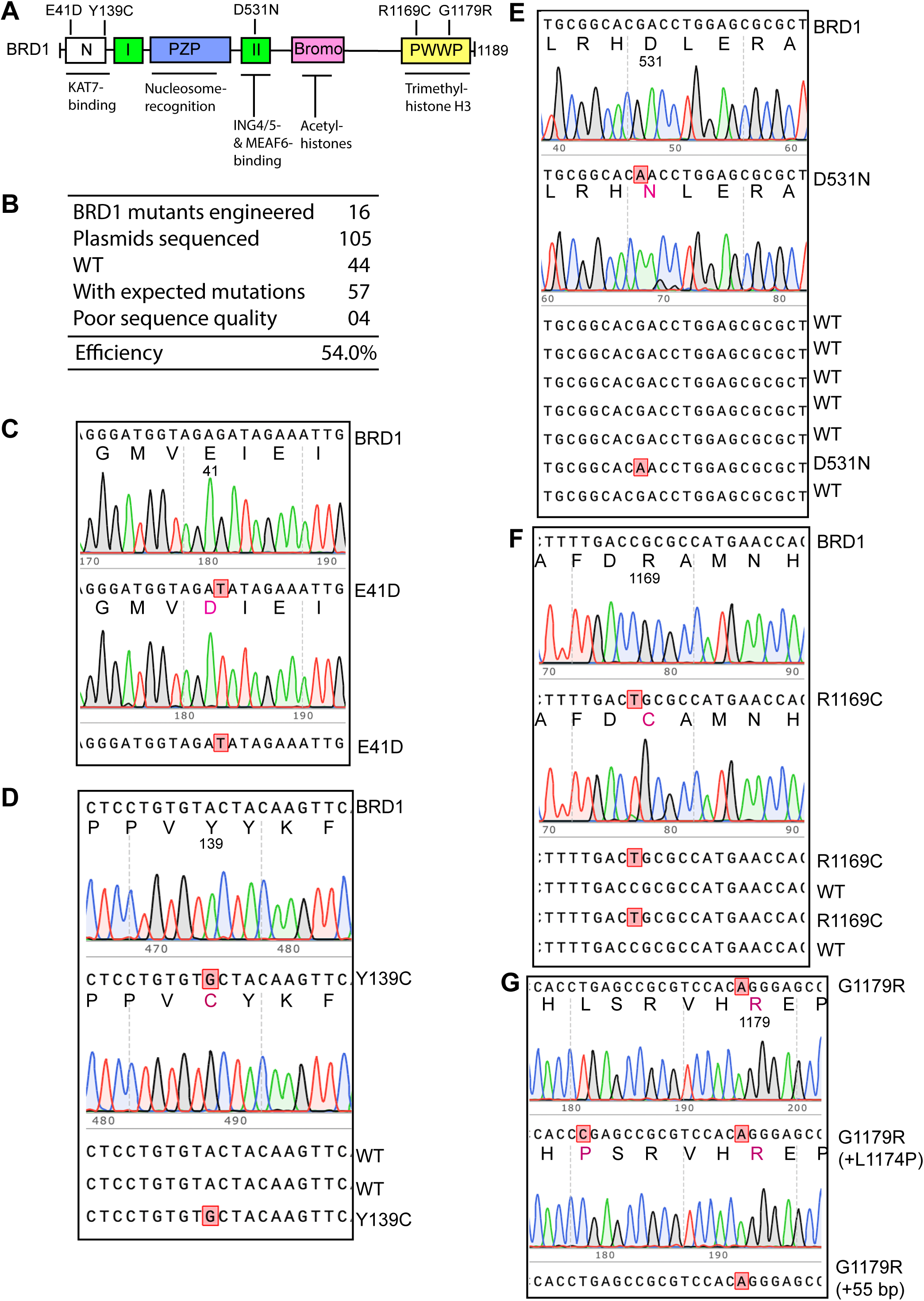
Efficient generation of BRPF2 mutants via P3 site-specific mutagenesis. A. Domain organization of BRPF2. It is composed of an N-terminal domain (N) required for interacting specifically with KAT7 [25], two EPC (Enhancer of Polycomb)-like motifs (I and II), a PHD-zinc buckle-PHD (PZP) module, a bromodomain and a PWWP domain [25]. For simplicity, only five missense variants are illustrated, with two located at the N-terminal domain (E41D and Y139C), one at the EPC-II motif (D531N) and two within the PWWP domain (R1169C and G1179R). B. Efficiency of the mutagenesis method. Among 105 plasmids sequenced, 4 could not be sequenced, perhaps due to wrong plasmids (such as the lack of primer sequence), poor plasmid quality or Sanger sequencing issues. To be conservative, they were all counted as incorrect mutant candidates for efficiency calculation. C. Sanger sequence analysis of 3 plasmids from engineering the E41D variant. Two were correct, so the mutagenesis efficiency was 66.7%. D. Sanger sequence analysis of 5 plasmids from generating the Y139C variant. Two were correct, so the efficiency was 40%. E. Sanger sequence analysis of 9 plasmids from engineering the D531N variant. Two carried the mutation, so the efficiency was 23%. F. Sanger sequence analysis of 6 plasmids from engineering the R1169C variant. Three were correct, so the efficiency was 50%. G. Sanger sequence analysis of 3 plasmids from engineering the G1179C variant. All three carried the mutation but two of them also habored unwanted insertions, so the efficiency was 33%. The mutagenesis reaction was carried out with Pfu-fly, which tends to introduce unwanted mutagenesis at the primer sites.

**Fig. 4.**
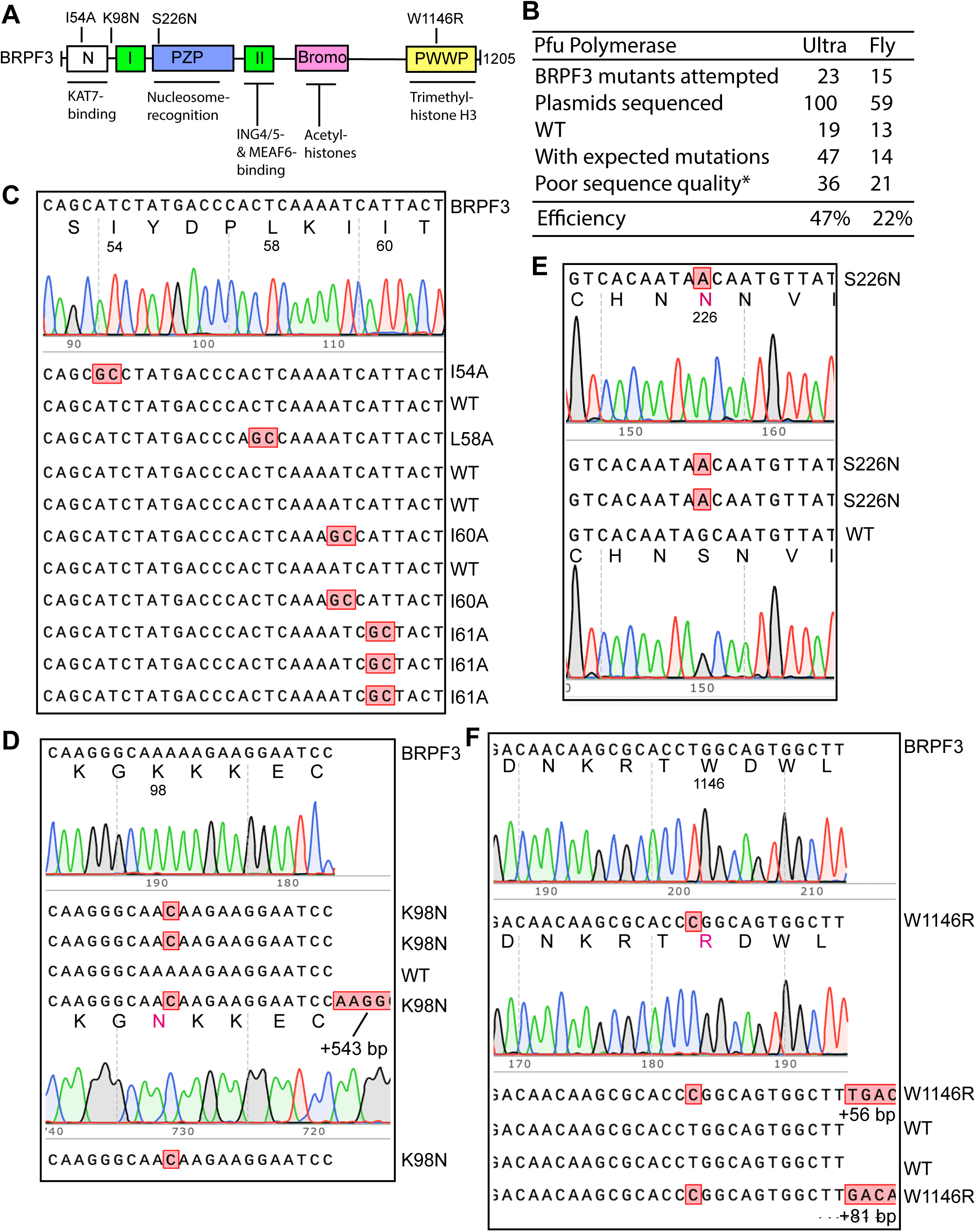
Efficiency of P3 site-directed mutagenesis to produce BRPF3 mutants. A. Domain organization of BRPF3. Like BRPF2, BRPF3 is composed of an N-terminal domain (N) for interacting with KAT7 [25], two EPC-like motifs (I and II), a PZP module, a bromodomain and a PWWP domain [25]. For simplicity, only four missense variants are illustrated, with two within or close to the N domain (I54A and K98N), one at the PZP module (S226N) and one at the PWWP domain (W1145R). B. Efficiency of the P3 mutagenesis method used for mutating a mammalian expression plasmid for HA-tagged BRPF3. Some mutagenesis reactions with PfuUltra yielded no or insufficient colonies for Sanger sequencing. By contrast, Pfu-fly polymerase rescued many of these failed experiments and generated sufficient colonies for Sanger sequencing, so despite the lower mutagenesis efficiency with Pfu-fly, it is still superior over PfuUltra in these cases. The asterisk denotes that plasmids with unwanted mutations were also included in the ‘poor sequence quality’ category. C. Sanger sequence analysis of 12 plasmids from engineering the I54A, L58A, I60A and I61A variant. For each, three plasmids were sequenced, with one (I54A or L58A), two (I60A) and three (I61A) being correct. Thus, the efficiency ranges from 33-100%. D. Sanger sequence analysis of 6 plasmids from engineering the K98N variant. Four contained the mutation but one of them possessed an insertion at the primer site, so the efficiency was 50%. E. Sanger sequence analysis of 4 plasmids from engineering the S226N variant. Three were correct, so the efficiency was 75%. F. Sanger sequence analysis of 6 plasmids from engineering the W1145R variant. Three contained the mutation but two of them also possessed insertions at the primer sites, so the efficiency was 16%.

According to the Clinvar database [1], there are also *BRPF3* mutations linked to genetic diseases. These mutations have not been molecularly characterized, so it is essential to engineer them and assess their pathogenicity. Moreover, for molecular characterization of the interaction of BRPF3 with KAT7, we needed to generate some artificial BRPF3 mutants. As the first step, we engineered some of these mutants (Fig. 4B-F). Unlike BRPF1 and BRPF2, we encountered difficulties in generating BRPF3 mutants. To obtain sufficient colonies for sequence analysis, 30 PCR amplification cycles were used instead of the standard 25 cycles. Despite this, there were still difficulties with some mutants. The main problem was that no or only few colonies could be obtained.

To overcome this, we tested PfuUltra II, a new version of PfuUltra with a 70-80% faster synthesis rate, but it yielded no colonies, so the effort was aborted. We also sought to replace PfuUltra with Pfu-fly as the latter was advertised to be a much faster enzyme and display much higher fidelity than PfuUltra (∼108 folds over Taq polymerase). Importantly, Pfu-fly rescued the situation. With the modified technique, we were able to engineer more BRPF3 mutants, at the efficiency varying from 16-100% (Fig. 4B-F). Some mutants (e.g. R15W) were still challenging to make. For this and also some of the BRPF3 mutants that we succeeded to engineer with Pfu-fly, we frequently detected unwanted deletions and/or insertions at the primer sites (Fig. 4D & 4F). Thus, further improvements are required to minimize unwanted mutagenesis and enhance reliability.

As for the underlying culprits that contribute to the difficulty with the human BRPF3 expression plasmid, one possibility is its special coding sequence. Related to this, its GC-content is one factor, reaching 90-95% in certain areas. For example, the GC-content for the coding sequence around R15 is close to 90% which may explain why R15W was difficult to engineer. On other hand, the GC-content of the coding sequence for BRD1 also reaches 90% is some areas, but it was OK to engineer its mutants. It is possible that the secondary or tertiary structures are also important factors.

### P3 mutagenesis to generate mutants of JADE and EPC epigenetic regulators

We have also extended the method to JADE2 and JADE3, two epigenetic regulators sharing several homologous domains with BRPF2 and BRPF3 [32]. Moreover, like them, JADE2 and JADE3 also interact with KAT7 to form highly stable complexes and activate their acetyltransferase activities [32]. Site-directed mutagenesis is needed to elucidate the molecular architecture of these acetyltransferase complexes. As the first step, we sought to engineer two JADE2 mutants, I76A and L77A, which are expected to affect the interaction with KAT7 [34,35]. Only one pair of primers was used. We sequenced 7 plasmids from the resulting colonies. Among them, five carried for the I76A substitution, one encoded L77A substitution and the 7th was a mixed clone of both mutants (Fig. S3A). For JADE3, we designed a truncation mutant, E463*, which is useful for assessing the function of the C-terminal portion of the protein. We analyzed three of the resulting colonies and found that all carried the expected E463* mutation (Fig. S3B). Thus, the mutagenesis efficiency to generate these three JADE2 and JADE3 mutants was 100%.

We have also tested the method with EPC1 and EPC2. As illustrated in Fig. S4A, they share two EPC-like modules with BRPF1, BRPF2 and BRPF3. Different from them, EPC1 and EPC2 serve as a scaffold to assemble tetrameric core acetyltransferase complexes with KAT5, ING3 and MEAF6 (Fig. S4A) [32]. The equivalent tetrameric complex in the budding yeast serves as the acetyltransferase core for the large NuA4 complex, whose structure and interaction with nucleosomes have been determined elegantly [36-39]. Moreover, structure of the mammalian TIP60 complex has also been solved [40,41]. Structure-based functional analysis should shed light on how EPC1 and EPC2 activate KAT5, which should also yield insights into how the related epigenetic regulators such as BRPF1 and its paralogs activate their respective acetyltransferases. As such, it is necessary to engineer EPC1 and EPC2 mutants. As the first step, we introduced stop codons and generated truncation mutants for EPC1 or EPC2 (Fig. S4A). As shown in Fig. S4B-F, the mutation efficiency was close to 75%. Thus, this method can be easily utilized to engineer EPC1 and EPC2 mutants.

### P3 mutagenesis for engineering different lysine acetyltransferase mutants

The expression plasmids for EPC1 and EPC2 are ∼7.8 kb in size, whereas the expression plasmids for BRPF1, BRPF2 and BRPRF3 are ∼9.2 kb. Thus, an interesting question is why it was less successful to generate BRPF3 mutants (Figs. 4 & S2). To investigate the general applicability of the method, we next tested it with four different mammalian expression vectors for FLAG-tagged lysine acetyltransferases, including KAT2B, KAT6A, KAT8, p300 and CBP. The size of these expression vectors ranges from 7.0-13.4 kb: human KAT8, 7.0 kb; KAT2B, ∼7.6 kb; KAT6A, 11.9 kb; p300, 12.7 kb and mouse CBP, 13.4 kb.

Some of the engineered mutations, including E570Q of KAT2B, K604R of KAT6A, K274R of KAT8, D1399Y/N of p300 and R1447C/H of CBP, were expected to inactivate the enzymes, so the resulting mutants serve as valuable negative controls for testing the enzymatic activities of these enzymes. For example, they are particularly useful for analyzing the newly discovered acyltransferase activities of these enzymes towards histones and non-histone proteins. Moreover, the p300 and CBP mutants are derived from cancer-associated hot-spot mutations, as reported in the COSMIC and cBioPortal databases (Fig. 5E) [2,3]. L1061* of KAT6A is derived from a patient with intellectual disability [16]. For D1399Y/N of p300 or R1447C/H of CBP, the primers were designed to engineer two different mutants using a single pair of primers with mismatch at the mutation sites, via the strategy illustrated in Fig. 1E-F. Notably, one pair of primers was successfully used to engineer two CBP mutants, Y1504D and Y1504H (Fig. 5D). This strategy was also utilized to generate the p300 mutants D1399Y and D1399N (Fig. 5F). As illustrated in Fig. 5A-D & 5F, the mutagenesis efficiency ranges from 75% to 100% for generating the KAT8, KAT6A, CBP and p300 mutants. Thus, the method should be useful for generating additional KAT8, KAT6A, CBP and p300 mutants.

**Fig. 5.**
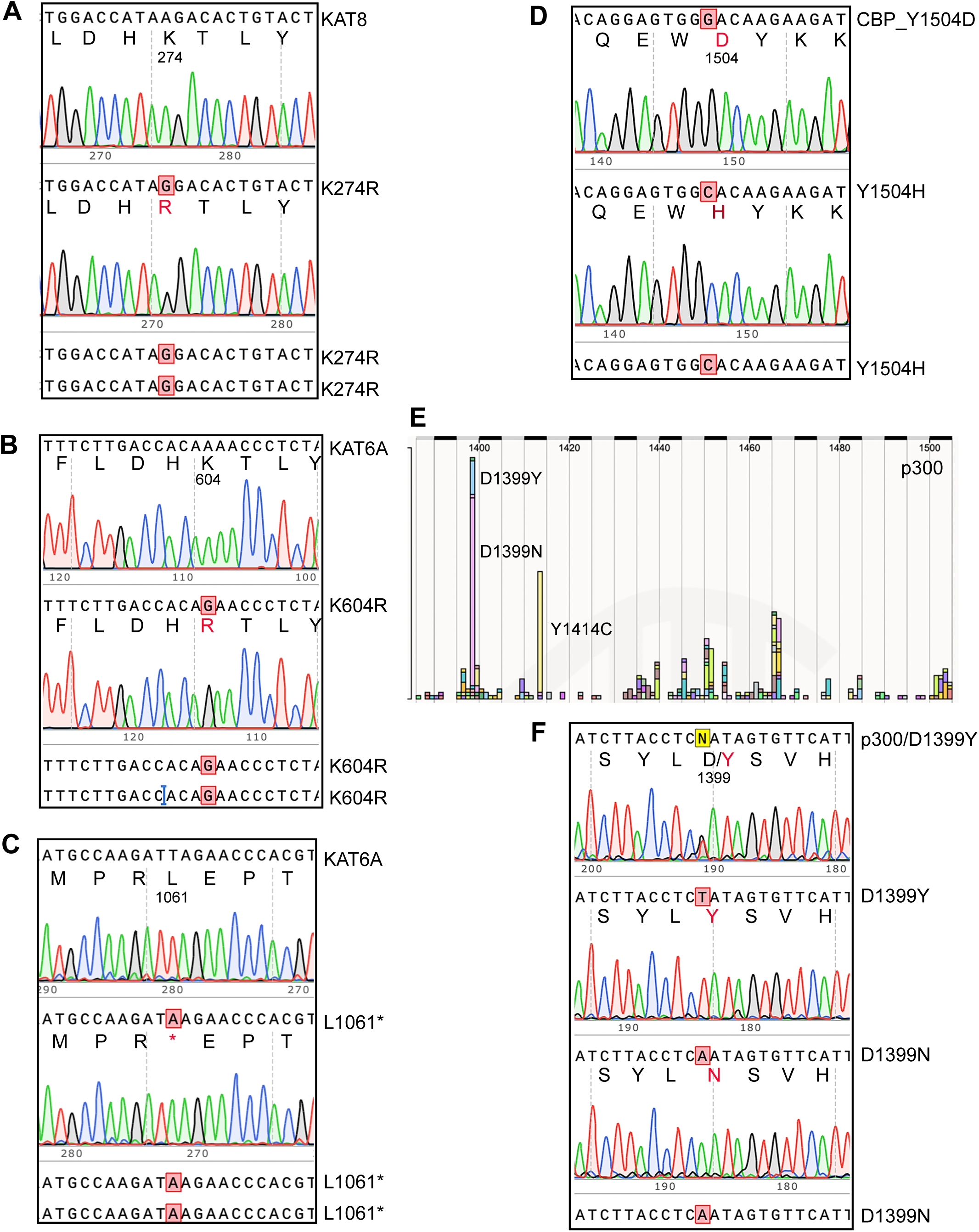
P3 mutagenesis for efficient construction of lysine acetyltransferase mutants. A. Sanger sequence analysis of 4 plasmids from engineering the K274R variant of KAT8. Three were correct, so the efficiency was 75%. B. Sanger sequence analysis of 4 plasmids from generating the K604R variant of KAT6A. Three were correct, so the efficiency was 75%. C. Sanger sequence analysis of 4 plasmids from engineering the K1061* variant of KAT6A. Three were correct, so the efficiency was 75%. D. Sanger sequence analysis of 4 plasmids from engineering the Y1504D and Y1504H variants of mouse CBP, equivalent to Y1503D and Y1503H of human CBP, two hot-spot mutations in different types of cancer. Among the 4 plasmids sequenced, one failed to be sequenced, one carried Y1504D and two had Y1504H, so the mutagenesis efficiency was 75%. Note that only one primer pair was used for the mutagenesis reaction, with the strategies depicted in Fig. 1E-F. E. The D1399Y and Y1399N mutants of p300 correspond to two hot-spot somatic mutations in cancer. The graph was copied from the COSMIC database [2,3]. F. Sanger sequence analysis of 4 plasmids from engineering the D1399Y and Y1399N variants of p300. Two harbored D1399Y and two carried Y1399N, so the mutagenesis efficiency was 100%. Note that only one primer pair was used for the mutagenesis reaction, with the strategies depicted in Fig. 1E-F.

One exception is the E570Q mutant of human KAT2B. Despite multiple trials with different PCR conditions, we failed to get any correct colonies. For each trial, the colony number was either low or none. Restriction digestion of plasmids from the few colonies obtained revealed large deletions, making us wonder whether the coding sequence of KAT2B is somewhat unique and thereby presents a barrier for PCR amplification. Related to this, during cloning and sequencing the corresponding cDNA [42], we noticed an extremely GC-rich region in the 0.27-kb coding sequence for the N-terminal 90 amino acid residues, reaching 95-100% in some areas. We tried to include 10% DMSO in the PCR buffer for PfuUltra and also tested Pfu-fly, but neither condition helped. As a result, we conclude that further improvements are needed for this and other plasmids with extremely GC-rich sequences.

### P3 site-directed mutagenesis for generating HDAC4 and HDAC5 mutants

In addition to lysine acetyltransferases and their regulators, we are interested in histone deacetylases, including HDAC4 and HDAC5 (Fig. S4A) [43]. A recent study has made a link of *HDAC4* mutations to a new developmental disorder [44], but the missense mutants have not been characterized adequately. For example, P248L occurs independently in three patients with intellectual disability [44], but it remains to be established whether this substitution is pathogenic and if so, how this substitution exerts its functional impact. A similar substitution, P248A, occurs in one patient with intellectual disability [44]. P248 is a critical residue for 14-3-3 binding [43], so we utilized P3 site-directed mutagenesis to engineer P248L and P248A by using a single pair of primers with mismatches at the mutation sites, via the strategy shown in Fig. 1E-F. We were able to obtain the mutant expression plasmids for GFP-tagged HDAC4 at a frequency of 60% (Fig. S5B). For FLAG-tagged HDAC4, we analyzed 4 plasmids. Among them, three were for P248L and one was for P248A, resulting in the ideal efficiency of 100%.

HDAC4 is paralogous to HDAC5 (Fig. S5A) [44]. They possess a serine-rich motif conserved from *Drosophila* to humans, but its functional significance remains unclear (Fig. S4A). To investigate this, we utilized P3 mutagenesis to generate mammalian expression plasmids for FLAG-tagged HDAC5 mutants S318A and S322A, at a frequency ranging from 25-30% (Fig. S5C-D). Thus, this method should be useful for generating additional HDAC4 and HDAC5 mutants.

### P3 mutagenesis for engineering missense spike variants of SARS-COV-2

The coronavirus disease 2019 (COVID-19) pandemic has resulted in tragic loss of life, affected health care systems and crippled the economy around the world. Severe acute respiratory syndrome coronavirus 2 (SARS-CoV-2) is the culprit behind this devastating disease [45]. Since its initial identification four years ago, it has gained various mutations and yielded many variants. Derived from Omicron variant, JN.1 and its descendants are contributing to active infections around the world. One major SARS-CoV-2 gene of variation encodes the spike protein, which is also the target of vaccines that have been or are being developed. Compared to Omicron variant, JN.1 and its descendants have gained additional mutations in the spike protein, including R346T, L455S, L456L, Q493E and V1104L [46]. To understand the underlying pathological mechanisms of action, it is necessary to engineer these substitutions in the parental spike protein for analysis of the functional impact. We thus utilized P3 mutagenesis for engineering these mutations in the spike protein backbones of D614G and Omicron variants. In addition, we engineered L981F in the backbone of spike protein of D614G variant. This substitution is present in Omicron variant. For L455S and L456L, we only utilized one pair of primers, with one encoding F456L and another encoding both L455S and F456L, via the strategies illustrated in Fig. 1E-F. As shown in Fig. 6A, the overall mutagenesis efficiency for these mutants is ∼50% when PfuUltra was used as the enzyme for PCR. Representative Sanger sequencing results are shown in Fig. 5B. These results indicate that this method is efficient and reliable for generating and studying new mutations that JN.1 and its decedents will continue to gain during evolution.

**Fig. 6.**
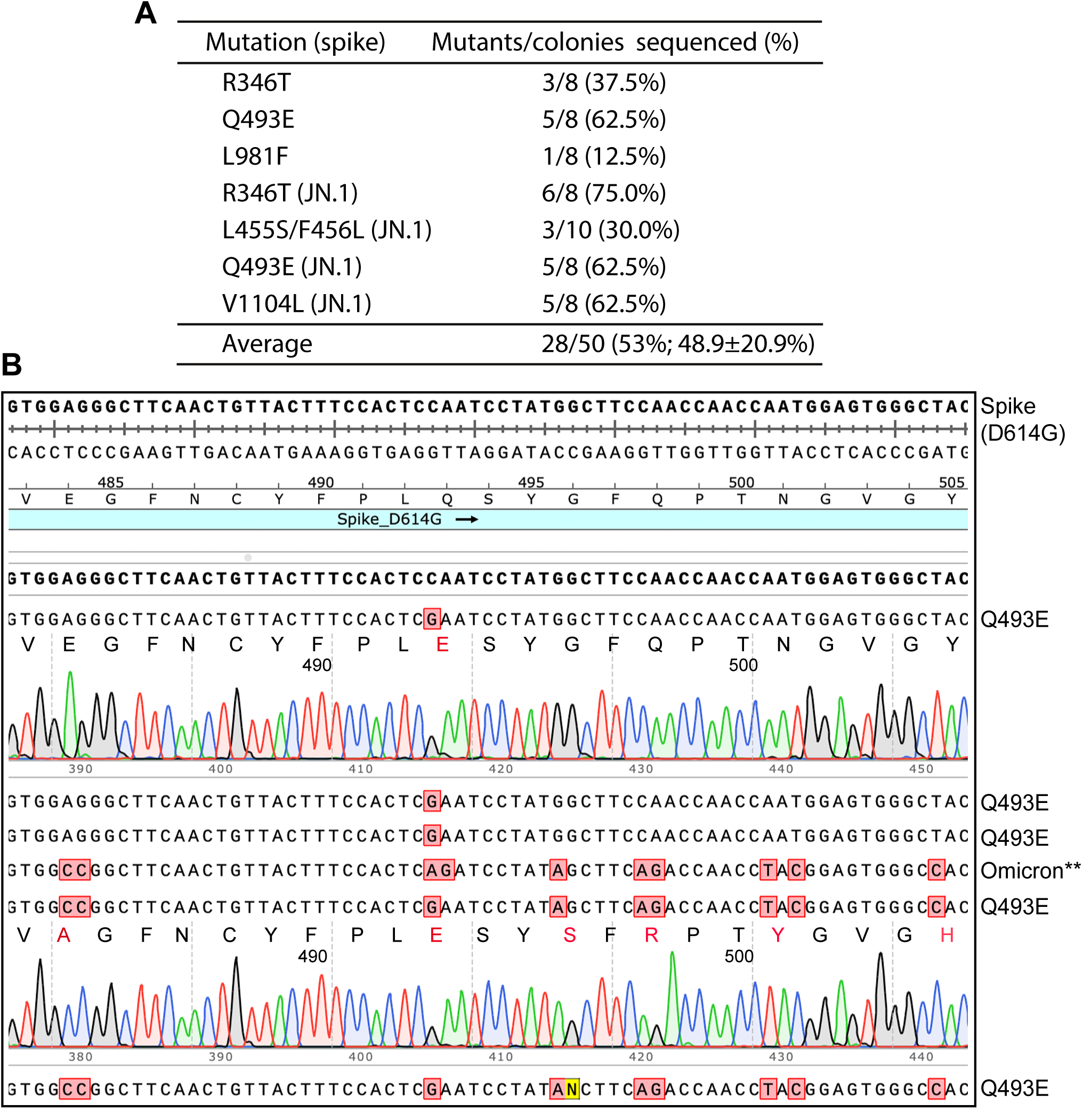
Efficient construction of SARS-COV-2 spike mutants via P3 mutagenesis. A. Efficiency for generating different spike mutants. The first three were generated in the backbone of D614G spike protein, whereas the last four were on the backbone of Omicron spike protein. The asterisk indicates that among the 10 colonies analyzed, three are mutants, with two for the L455S/F456L mutant and one for F456L. For the average efficiency, standard deviation is also provided. For L455S/F456L and F456L mutants, we used a single pair of primers. The mismatch between the two primers at the overlapping region may be one factor contributing to the low efficiency. Related to this, Pfu-Fly performed much better than PfuUltra used for this panel (see Fig. S6). As for L981F, it remains unclear, but one possibility is that the quality of the primers, but for over a hundred of mutations that we have engineered, such a low efficiency is an extremely rare event. B. Sanger sequence analysis of representative plasmids from engineering the *SARS-COV-2* spike mutation Q493E. The top three candidates were from the mutagenesis experiment with the expression plasmid for the D614G spike protein as the template, whereas for the bottom three the expression plasmid for the Omicron spike protein as the template. In addition, Q493E, other mutations that Omicron carries in the surrounding regions are also boxed in red. In Omicron variant, Q493 is replaced with arginine.

As stated above, Pfu-fly polymerase shows higher fidelity and requires less PCR amplification time than PfuUltra. We thus utilized Pfu-fly to generate three spike mutants. For engineering the F456L mutant and the L455S and F456L double mutant, a single pair of primers was used. As shown in Fig. S6A, we analyzed 4 colonies and found that three encoded the double mutant and one was for the F456L mutant. Thus, the mutagenesis efficiency was 100%. For the V1104L mutant, we analyzed four colonies and found that three of them carried the mutant plasmid (Fig. S6B), so the mutagenesis efficiency was 75%. These results indicate that this method is highly efficient for generating the three new mutations that JN.1 and its decedents have gained during evolution.

## Discussion

Based on PCR with completely complementary primer pairs (Fig. 1A), the QuickChange^™^ site-directed mutagenesis method has been widely used [9,10]. An alternative approach is to utilize a pair of partially complementary primers with two 3’-protruding ends (Fig. 1B), thereby overcoming limitations intrinsic to the QuickChange^™^ site-directed mutagenesis method (Fig. S1A) [12,13]. As the method was developed and tested primarily for 5-6 kb plasmids [13], it remains unclear how it performs with larger plasmids and those with more complex sequences, as site-directed mutagenesis typically becomes more challenging with larger plasmids [18-20]. In this regard, we have systematically evaluated the newly optimized method (Fig. 1D-F) with dozens of mutations in a dozen of mammalian expression vectors. The size of the expression vectors we tested ranges from 7.0-13.4 kb, and many of them are much larger than the 5-6 kb plasmids tested previously [13]. The new method was successfully adapted to generate mutants for over a dozen of mammalian expression plasmids, including those for BRPF1 (Fig. 2), BRPF2 (Fig. 3), JADE2 (Fig. S3), JADE3 (Fig. S3), EPC1 (Fig. S4), EPC2 (Fig. S4), KAT6A (Fig. 5B-C), KAT8 (Fig. 5A), CBP (Fig. 5D), p300 (Fig. 5E), HDAC4 (Fig. S5), HDAC5 (Fig. S5) and the spike protein of SARS-COV-2 (Figs. 6 & S6). The average efficiency was ∼50%, but the mutagenesis efficiency reached 100% in some cases, such as KAT8 (Fig. 5A), JADE2, (Fig. S3A), JADE3 (Fig. S3B), HDAC4 (Fig. S5 and the spike proteins of two SARS-COV2 variants (Figs. 6A & S6). This is not something that we have observed with the QuickChange method [14-17]. Moreover, the QuickChange method tends to become more problematic with large plasmids that reach over 10 kb. By contrast, this is not issue with P3 mutagenesis.

Intriguingly, it is more challenging to generate BRPF3 mutants with this new method (Fig. 3). An important question is why this expression vector is more difficult than the others. One possibility is its relatively high GC-content, but it is very similar to BRPF2 in this regard. Alternatively, it is because of the presence of some special or complex sequences or even unique secondary structures. Related to this, certain sequences inhibit proofreading of DNA polymerases [47]. Importantly, Pfu-fly appeared to alleviate the problem with BRPF3 to certain extent (Figs. 4B & S2) although it still failed to engineer some mutants, such as R15W (Fig. S2A). A problematic and puzzling issue is that this polymerase introduces more unwanted deletions and insertions at the mutation or primer sites (Figs. 4 & S2A), which might have contributed to the significantly lower mutagenesis efficiency observed (Fig. 4B). Intriguingly, the KAT2B plasmid was even more challenging, perhaps due to an extremely GC-rich region [42]. Thus, while the newly optimized mutagenesis method is efficient for engineering many mutations, further improvement is still needed to make it more reliable and efficient for some cases.

In some experiments, we utilized a single primer pair to introduce two mutations at neighboring sites (Fig. 1D-F). Thus, an interesting issue is whether two or more distant mutations can be engineered in the same mutagenesis reaction Fig. S1). This was explored in an earlier study [13]. Theoretically, the amplification efficiency of internal PCR products from primers at two or more mutation sites is significantly higher than amplification of the entire plasmid templates (Fig. S1C-D). This disparity makes it challenging to adapt the method for simultaneously engineering two or more distant mutations. To overcome this limitation, one potential solution is to carry out the multiple mutation reactions sequentially. Alternatively, a new method needs to be developed for simultaneous multi-site mutagenesis, which would be time saving in comparison to the sequential approach.

One common issue with site-directed mutagenesis is potential unwanted mutations. There mutations can arise from several sources: 1) impurity in primers, 2) unwanted recombination at the mutation or primer sites, and 3) misincorporation into newly synthesized strands introduced by DNA polymerases. Related to source 3), PfuUltra DNA polymerase is a high-fidelity enzyme, 3-fold higher than Pfu itself [23]. Based on our experience with hundreds of Sanger sequencing reactions, misincorporation during synthesis mediated by DNA polymerase is extremely rare. By contrast, both sources 1) and 2) are much more significant. To minimize unwanted mutations due to impurity in primers, one approach is to use acrylamide-purified oligonucleotides. However, this method is more time-consuming and significantly more expensive than using standard desalted oligonucleotides. Alternatively, reducing the length of oligonucleotides can also mitigate the risk of unwanted mutations. Related to this, we have utilized primers ∼30 nucleotides in length, significantly shorter than the previous studies [12,13]. Remarkably, even when using just desalted nucleotides for mutagenesis, unwanted mutations at the primer sites are rare events in experiments based on PfuUltra, and any such mutations can be easily identified during Sanger sequencing. This highlights the practicality of our approach while maintaining the integrity of the mutagenesis process.

As for unwanted recombination at the mutation or primer sites, it has been observed in a few cases (Figs. 2E, 3G, 4D, 4F & S2). The frequency increased as the PCR cycle number rose from 25 to 30 cycles (Fig. 4) or when Pfu-fly was used (Figs. 3G & S2). This decreased the mutagenesis efficiency to some extent. One reason for such unwanted recombination events is the strand-displacing activity and/or low processivity of the DNA polymerases that were used. This also reflects how faithfully the strand synthesis stops when reaching the 5’-end of an annealed primer. To minimize the impact, we carried out some PCR reactions at 68°C, instead of 72°C [12,13]. But this change was unable to address the issue, so further studies are needed to resolve this important issue. Contrastingly, this issue did not appear to be a problem with the two plasmids for the spike protein of SARS-COV-2. With those plasmids, both PfuUltra and Pfu-fly were highly efficient (Figs 6 & S5). Impressively, with Pfu-fly, three spike mutants were generated within ∼3 days at the ideal efficiency of 100% (Fig. S5). The different efficiency of Pfu-fly with the expression vectors for BRPF3 (Figs. 4B & S2) and the SARS-COV-2 pike protein (Fig. S5) suggests that plasmid sequences may be an important determinant for mutagenesis efficacy. In addition to engineer mutants of the spike protein for functional assays and vaccine development, the current method should be applicable to antibody engineering as required for developing therapeutics to fight against SARS-COV-2.

One unexpected finding is that PfuUltra II did not work at all, although the underlying reason remains unclear. Performance of three different Pfu derivatives (i.e. PfuUltra, PfuUltra II and Pfu-Fly) also suggests that the quality (including fidelity, processivity and synthesis rate) of the thermostable DNA polymerase used for PCR is highly important for the success of the mutagenesis experiments. To improve this method further (e.g., to reach the ideal efficiency of 100%), one important direction is to test additional thermostable DNA polymerases. Related to this, there are new thermostable DNA polymerases that are reported to be superior to Pfu. It is likely that some of the enzymes will make the current method much better. Another direction is to figure out ways by which plasmids with extremely GC-rich sequences (such as the KAT2B expression vector used here) can be easily used for site-directed mutagenesis. As many proteins contain domains with their corresponding coding sequences are GC-rich, so this research direction shall be important for a site-directed mutagenesis generally applicable. Thus, the current study also provides a simple conceptual framework on how to develop the method further.

In summary, we have optimized a new site-direct mutagenesis method (Fig. 1D-E) and systematically evaluated it with >100 mutations on a dozen of mammalian expression vectors, ranging from 7.0-13.4 kb. Compared to the QuickChange^™^ mutagenesis method, the success rate has increased significantly, reaching an average efficiency of ∼50%, at or close to 100% in some cases, and often requiring sequence analysis of only 2-3 bacterial colonies per mutation reaction. Notably, Pfu-fly is superior to PfuUltra in certain cases (Figs. 4, 6, S2 & S5). Results from this study also shed light on how to improve the method further by enhancing its efficiency and reliability. Interestingly, among over a dozen of expression plasmids that we have tested, the one for BRPF3 is more difficult to mutate than the others (Figs 4 & S2), perhaps due to its unique sequence. Therefore, while there is still room for refinement, this newly optimized mutagenesis method represents a significant advancement, being efficient, economical and versatile across different expression vectors. It effectively eliminates site-directed mutagenesis as a bottleneck in basic and clinical research. Results from this study also sheds light on further improvements required for setting up ideal site-specific mutagenesis methods with the ideal efficiency at or close to 100% to mutate a wide spectrum of plasmids rapidly and reliably.

## Experimental Procedures

### Plasmid vectors

Mammalian expression vectors for HA-tagged BRPF1, BRPF2 (also known as BRD1) and BRPF3 were described earlier [16,31,48]. The mammalian expression vector for FLAG-KAT6A was kindly provided by Dr. Issay Kitabayashi [49]. The expression plasmids for FLAG- or GFP-tagged HDAC4 were reported previously [50,51]. The expression plasmids for FLAG-tagged HDAC5, untagged p300 and FLAG/HA-tagged mouse CBP were obtained from Addgene (Cat. 32213, 23252 and 32908, respectively). The coding sequence for the FLAG tag was engineered into the p300 expression vector and the coding sequence for the HA tag was deleted to express FLAG-tagged p300 and CBP, respectively, with the details to be published elsewhere. The expression plasmid for FLAG-tagged human KAT2B (a.k.a. PCAF) was reported previously and is similar to the one available from Addgene (Cat. 8941, but with a different promoter for mammalian expression [42]). The expression plasmids for HA-tagged EPC1 and EPC2 were constructed from a pcDNA3.1 derivative that encodes an HA tag, with the details to be published elsewhere. The mammalian expression vectors for untagged D614G and Omicron spike proteins of SARS-COV-2 were purchased from SinoBiological (Cat. VG40589-UT and VG40835-UT, respectively).

### Mutagenesis primers

Primers were designed with the aid of SnapGene software package. They were synthesized at Integrated DNA Technologies (IDT) in the 25-nmol scale as standard and desalted DNA oligos, without further purification. Representative primers utilized to engineer BRPF1 mutants were as listed below, where the mutated nucleotides are shown in the lower case: P19S-F, GCGACTAAGtCACCATACGAGTGCCCGGTGGA; P19S-R, CGTATGGTGaCTTAGTCGCCCGCAAGTTGTGG; P19S-F1, GGCGACTAAGtCACCATACGA; P19S-R1, TCGTATGGTGaCTTAGTCGCC; C23R-F, CCATACGAGcGCCCGGTGGAGACCTGCCGAAAG; C23R-R, CCACCGGGCgCTCGTATGGTGGCTTAGTCGCCC; C23R-F1, ACCATACGAGcGCCCGGTGGA; C23R-R1, TCCACCGGGCgCTCGTATGGT; R66C-F1, AAAAAGGGGtGCCAGTCACGCCCAGCCAAC; R66C-R1, GTGACTGGCaCCCCTTTTTCTTGTGCTTGC; F1154del-F, TCGTCCTCTTCGACAACAAACGAACCTGGCAGTGGCTGCCCAGGA; and F1154del-R, CGTTTGTTGTCGAAGAGGACGAGGTAGAGATGCTCTCGGGCTTCC. P19S-F1, P19S-R1, C23R-F1 and C23R-R1 were designed based on the strategy shown in Fig. 1A.

Primers utilized to generate BRD1 mutants were as follows: E41D-F, GGATGGTAGAtATAGAAATTGAAGGGCGCT; E41D-R, AATTTCTATaTCTACCATCCTTTGAGCTTG; Y139C-F, CTCCTGTGTgCTACAAGTTCATCGAGAAGT; Y139C-R, AACTTGTAGcACACAGGAGGCCTCCTGGGG, D531N-F, CTGCGGCACaACCTGGAGCGCGCTCGCCTG; D531N-R, GCTCCAGGTtGTGCCGCAGCCGCTGCCAGT, E41D-F1, GGATGGTAGAtATAGAAATTG; E41D-R1, CAATTTCTATaTCTACCATCC; Y139C-F1, CCTCCTGTGTgCTACAAGTTC; Y139C-R1, GAACTTGTAGcACACAGGAGG; E1107Q-F, AAGATTGGGcAGCACATGCAGACCAAGTCT; E1107Q-R, GCATGTGCTgCCCAATCTTCAGCACGTCCA; I1146V-F, GACGAAACTgTAGACAAGTTAAAGATGATG; I1146V-R, ACTTGTCTAcAGTTTCGTCAATACCAAGGG; G1179R-F, CGCGTCCACaGGGAGCCGACCAGCGACCTC; R1169C-F, GCTTTTGACtGCGCCATGAACCACCTGAGC; R1169C-R, TTCATGGCGCaGTCAAAAGCGATCCGCACG and G1179R-R, TCGGCTCCCtGTGGACGCGGCTCAGGTGGT.

Primers utilized for engineering BRPF3 mutants were I52A-F, GCCTGCATCGTgcCAGCATCTATGACCCACT; I52A-R, TAGATGCTGgcACGATGCAGGCGTCCATCAA; I54A-F, CGTATCAGCgcCTATGACCCACTCAAAATCA; I54A-R, GGGTCATAGgcGCTGATACGATGCAGGCGTC; L58A-F, TATGACCCAgcCAAAATCATTACTGAAGATG; L58A-R, ATGATTTTGgcTGGGTCATAGATGCTGATAC; I60A-F, CCACTCAAAgcCATTACTGAAGATGAGCTAA; I60A-R, TCAGTAATGgcTTTGAGTGGGTCATAGATGC; I61A-F, CTCAAAATCgcTACTGAAGATGAGCTAACTG; I61A-R, TCTTCAGTAgcGATTTTGAGTGGGTCATAGA; K98N-F, CAAGGGCAAcAAGAAGGAATCCTGCTCCAA; K98N-R, TTCCTTCTTgTTGCCCTTGGATGAGGGTTT; S226N-F, GTCACAATAaCAATGTTATTCTCTTCTGTG; S226N-R, ATAACATTGtTATTGTGACATTCATCATCC; K1075E-F, GGAGACCTGGAGCCCTTGGAGCTGGTGTGG; K1075E-R, CCAAGGGCTCCAGGTCTCCGCGGTCTTCAA; W1145R-F, AAGCGCACCaGGCAGTGGCTTCCAAGGGAC; and W1145R-R, GCCACTGCCtGGTGCGCTTGTTGTCAAAGA. The primers K1075E-F and K1075E-R were to repair an unexpected mutation in the mammalian expression plasmid for HA-tagged expression plasmid.

The primers for JADE2 were as follows: I76A-F, TGACTACTACgctCTGGCAGACCCATGGCGACAGGA; and L77A-R, ATGGGTCTGCagcGATGTAGTAGTCATCCGGGCTGA. The primers for JADE2 were as follows: E463*-F, CCAAAAGAGtGAAAGCATTCACACTCGAAT; E463*-R, AATGCTTTCaCTCTTTTGGCTGCACCAGCC.

The primers for KAT8 were as follows: K274R-F2, TGGACCATAgGACACTGTACTTTGACGTGG; and K274R-R2, TACAGTGTCcTATGGTCCAGGAAAAGCTTG. The primers for engineering KAT2B mutants were as follows: E570Q-F, GGATTCACAcAGATTGTCTTCTGTGCTGTA; and E570Q-R, AGACAATCTgTGTGAATCCTTGAGATGGGA. The primers for KAT6A were K604R-F, TTGACCACAgAACCCTCTATTACGATGTGG; K604R-R, TAGAGGGTTcTGTGGTCAAGAAACAACTTT; L1061*-F, ATGCCAAGATaAGAACCCACGTTTGAGATCGATGAAGAAGAGGAG; and L1061*-R CGTGGGTTCTtATCTTGGCATTGGCCTCTCGGAGTCAGAATCTTC. The primers for p300 were D1399Y-F, ATCTTACCTCtATAGTGTTCATTTCTTCCGT; and D1399N-R, ATGAACACTATtGAGGTAAGATATGTATACT. The primers for mouse CBP were as follows: R1446C-F, CCGCTGCCTCtgtACAGCTGTTTACCATGAGAT; R1446H-R, AACAGCTGTatgGAGGCAGCGGGGCCGGAAGA; Y1503D-F, ACAGGAGTGGgACAAGAAGATGCTGGACAAG; and Y1503H-R ATCTTCTTGTgCCACTCCTGTAGTCGTTTTG.

Primers for engineering HDAC4 mutants were P248L-F, GCTTCTGAACtGAATCTGAAATTACGGTCCA; and P248A-R, TTCAGATTCGcTTCAGAAGCTGTTTTCCTAA. Primers for HDAC5 were as follows: S318A-F, GCACCCGGCgctGGCCCCAGCTCTCCCAACAG; S318A-R GCTGGGGCCagcGCCGGGTGCGCTGTTACACA; S321A-F, CGGCCCCgctTCTCCCAACAGCTCCCACAG, S321A-R, TGGGAGAagcGGGGCCGGAGCCGGGTGCGC; S322A-F, GGCCCCAGCgCTCCCAACAGCTCCCACAGC; and S322A-R, TGTTGGGAGcGCTGGGGCCGGAGCCGGGTG.

Primers for engineering mutations into two expression vectors for the spike protein of SARS-COV-2 were R346T-F, ATGCCACCAcGTTTGCCTCTGTCTATGCCT; R346T-R, GAGGCAAACgTGGTGGCATTGAACACCTCT; F456L-F, TACAGACTGcTCAGGAAGAGCAACCTGAAA; L455S-R, GCTCTTCCTGAggctTCTGTAGAGGTAGTTGTAG; Q493E-F, TTTCCACTCgAATCCTATGGCTTCCAACCA; Q493E-R, CATAGGATTcGAGTGGAAAGTAACAGTTGA; Q493E-F1, CTTTCCACTCgaATCCTATAGCTTCAGACCAA; Q493E-R1, GCTATAGGATtcGAGTGGAAAGTAACAGTTGA; L981F-F GAATGACATCTTCAGCAGACTGGACAAGGTGGA, L981F-R, CCAGTCTGCTGAAGATGTCATTCAGCACAGAGG; V1104L-F, CACTGGTTTcTGACCCAGAGGAACTTCTAT; and V1104L-R, TCTGGGTCAgAAACCAGTGGGTGCCATTGC. Notably, at its 3’-end, the R346T-R primer has a mismatch with the coding sequence for Omicron spike protein; perhaps due to 3’→5’ exonuclease activity of Pfu, the primer could be still used even for this DNA template.

### Site-directed mutagenesis via PfuUltra high-fidelity DNA polymerase

For a mutagenesis reaction, the template plasmid was isolated from DH5α (could be any other Dam^+^ *E. coli* hosts), which is necessary for subsequent DpnI digestion. Primers were designed with the aid of the SnapGene software package (version 7.2.1) and purchased from IDT as small-scale desalted oligos, without further purification. Their sequences were listed above. If primer sequences contain long stretches of A and/or G (typically more than 5), a silent mutation was introduced to break such stretches. This was also done with long stretches of T or C (typically more than 5). All oligonucleotides were used without polyacrylamide gel or HPLC purification. Upon receipt from IDT, autoclaved Nanopure water, from an Atrium Pro water purification system (Satorius), was used to dissolve lyophilized oligonucleotides to prepare 100 mM stocks for further dilution to 10 mM working solutions.

PCR reactions were set up in 0.2 ml 8-strip thin-wall PCR tubes (Diamed, Cat. DIATEC420-1378), with each reaction containing 10-15 ng of plasmid DNA, 100 μM dNTPs, 0.2 mM forward primer, 0.2 mM reverse primer, 1x Pfu reaction buffer and 1.25 units of PfuUltra (Agilent, Cat. 600380) or PfuUltra II Fusion (Agilent, Cat. 600670) high-fidelity DNA Polymerase in autoclaved Nanopure water in a final volume of 10 μl. Amplification was carried out in a Bio-Rad PCR T100 Thermal Cycler using the following parameters: 95-96°C for 2-3 min as the initial step to denature plasmid DNA, followed by 20-30 cycles to amplify the DNA. Each amplification cycle was composed of 92-93°C for 15 seconds as the denaturation step, 50-52°C for 15-20 seconds as the annealing step and 68°C (or 72°C) for 6-9 minutes as the extension step. The extension time was calculated based on the plasmid size, with one minute allocated per kilobases (kb). The amplification cycle number varied from one plasmid to another, with the initial number set at 20; if it did not work, it was increased to 25-30. After the amplification step, an addition extension step of 68°C (or 72°C) for 10-15 minutes was added to the end of the cycling program. After that, the reaction was pre-programmed for short-term storage at 4°C, if the reaction was not digested immediately by DpnI (New England Biolabs, Cat. R0176). For DpnI digestion, 0.25 μl (20 units/μl) was added directly to the PCR reaction mixture (without extra buffers), which was then pipetted into a clean PCR tube for further incubation at 37°C for 90 min in the BioRad T100 Thermal Cycler.

According to the manufacturer’s instructions, PfuUltra II Fusion DNA polymerase was supposed to be superior to PfuUltra DNA polymerase. However, due to some unknown reasons, amplification with PfuUltra II Fusion DNA polymerase yielded no or few colonies, so this enzyme was not pursued further in this study.

### Site-directed mutagenesis via TransStart® Pfu-fly DNA polymerase

Per the manufacturer’s instruction, PfuUltra synthesizes at the rate of about 1 kb per minute, resulting in prolonged PCR time for large plasmids and potentially decreasing overall efficiency. Thus, it is beneficial to speed up the synthesis rate and reduce this extension step during PCR. To address this, TransStart® FastPfu Fly DNA polymerase (referred to as Pfu-fly herein) was developed. Therefore, it was purchased (GeneBio Systems Inc™, Cat. AP231-01) and tested for engineering BRPF3 mutants, some of which were difficult to obtain when PfuUltra DNA polymerase (Agilent, Cat. 600380) was used. The PCR parameters were the same as described for above PfuUltra DNA polymerase, except that the extension time for PCR was reduced from 1 min to 30 seconds per kb of a plasmid template. As a result, the PCR amplification time was shortened substantially. Moreover, compared to PfuUltra DNA polymerase, TransStart® FastPfu Fly DNA polymerase yielded ∼5 times more colonies for many BRPF3 mutagenesis reactions. As a result, the reaction volume could be reduced further to 5 μl, which led to the minimal reagent cost of $0.5 per mutagenesis reaction.

### Transformation of DH5α competent cells

To minimize microbial contamination, the following step was carried out near a gas flame from a Bunsen burner. 2-5 μl of the DpnI-digested PCR mixture was gently mixed with 20-50 μl of DH5α competent cells in a pre-chilled 0.5 ml sterile Eppendorf tube and incubated on ice for 30 min. For mutagenesis with Pfu-fly, 2 μl of the digested PCR mixture generated sufficient colonies, but 5 μl was required for a mutagenesis reaction based on PfuUltra. The remaining digested PCR mixture was frozen at -20°C as a backup. After the 30-min incubation on ice, the tube containing the DH5α competent cells and DpnI-digested PCR mixture was subjected to heat-shock at 42°C for 45 seconds and immediately returned to ice. About the length of heat shock, we also tested 30 seconds and found that heat-shock for 45 seconds yielded 10-15% more colonies. 50-125 μl of cold SOC (<u>s</u>uper <u>o</u>ptimal broth with <u>c</u>atabolite repression) medium was then added to the tube and after gentle mixing by tapping, the tube was incubated at a 37°C water bath for 30-45 min, without shaking. Afterwards, the entire cell mixture was pipetted onto a LB-agar plate as 4-5 large drops; the plate contained 100 μg/ml ampicillin (see the recipe below) or another antibiotic as required for the plasmid. An autoclaved 1.5 ml Eppendorf tube could be held on its cap allowing the ventral part of the lower half to be used for spreading out the cell mixture evenly on the plate. Afterwards, the plate was inverted and placed in a 37°C incubator (humidified with a beaker or flask of deionized water) for 18-24 h to allow bacterial colonies to appear. To reduce the usage of LB-agar plates, one transformation mixture could be plated onto one half of an LB-agar plate containing ampicillin or another proper antibiotic, allowing two transformation mixtures onto a single plate; to avoid cross-contamination, the mixtures were plated onto areas away from the central line.

With a sterile 200 μl pipette tip, a single colony was inoculated into a tube containing 6 ml LB supplemented with ampicillin (100 μg/ml) or another antibiotic as required for the plasmid. Typically, 2-3 colonies were initially analyzed per mutation; if none was the mutant, additional colonies were analyzed. When more than one colony was analyzed, special care was taken to avoid erroneous mixing of the corresponding tubes. For convenience, 100 mg/ml ampicillin aliquots were prepared as follows: 0.5 g ampicillin powder was dissolved with 5 ml of autoclaved Nanopure water in a 15 ml Falcon tube, separated into five 1.5 ml sterile Eppendorf tubes and kept at -20°C. If a clean spatula was used when ampicillin powder was weighted, no filtration of its solution was necessary.

After incubation in a bacterial shaker for 18-24 h, 0.5 ml of the bacterial culture was mixed with 0.25-0.5 ml 50% glycerol for long-term storage at -80°C. With a Qiagen miniprep kit (Cat. 27014), the plasmid was isolated from the remaining 5.5 ml bacterial culture. Plasmid was eluted in 50-150 μl autoclaved Nanopure water and 1μl was used to measure the concentration on a Nanodrop UV-Visible Spectrophotometer (Thermo Scientific, model 200). The plasmid was subsequently subject to Sanger sequencing. The resulting DNA sequences were compared with the wild-type template for the presence of the desired mutations using the sequence alignment tools in the SnapGene software package. The corresponding chromatograms were manually inspected for sequencing quality. The resulting DNA sequences and their corresponding chromatograms were also examined for potential unwanted mutations. Typically, only three colonies were needed to be analyzed to obtain a mutant.

### Preparation of LB (<u>L</u>uria-<u>B</u>ertani) liquid medium and LB agar plates

To prepare LB medium, 10 g tryptone, 5 g yeast extract and 5 g NaCl were added into a 1-liter glass bottle with a screw cap. 970 ml deionized or double distilled water was added to the bottle to make the total volume of 1 liter, along with 1 ml of 2 M NaOH to adjust PH to ∼7.0. With the screw cap being loosened slightly, the bottle was sterilized by autoclaving for 20 min at 15 psi (1.05 kg/cm^2^) on a liquid cycle. After cooling down, the cap was tightened for storage at room temperature; once the bottle was opened and recapped, it was kept at 4°C for long-term storage for up to a few months.

To prepare LB agar plates, 10 g tryptone, 5 g yeast extract, 5 g NaCl and 15 g agar were weighted directly into a 2-liter Erlenmeyer glass flask. 970 ml deionized or double distilled water was added to the flask to make the total volume of 1 liter, along with 1 ml 1 M NaOH to adjust pH ∼7.0. After the opening was covered with a sheet of aluminium foil, the flask was sterilized by autoclaving at 15 psi (1.05 kg/cm^2^) for 20 min on a liquid cycle. After sterilization, the flask was transferred to a 50-52°C water bath to cool down. While waiting for the medium to cool down to 50-52°C, 40-50 sterile petri dishes were taken out from their original packing plastic bags (Fisher FB0875712), which were cut open only from one end so that they would be used later for storage of the LB-agar plates (see below).

Once the medium was cooled down to the water bath temperature (requiring 0.5-1 h), 1 ml 100 mg/ml ampicillin (or another antibiotic as required for the plasmid) was added into the medium and the flask was gently swirled to mix ampicillin thoroughly with the medium, without generating too many air bubbles. The flask was transferred to a bucket containing some 50-52°C water and moved to a bench. Near a gas flame, the molten LB-agar-ampicillin medium was pipetted, with a 25 ml sterile plastic pipette, into the sterile petri dishes at 20 ml per dish. To avoid premature gelling, this step was done as swiftly as possible. If needed, air bubbles in the gelling LB-agar plates could be killed carefully with the flame. The plates were stacked at 5-10 per deck for complete gelling on the bench. After being kept on the bench overnight (for agar surface drying), the plates were put back into the original plastic packing bags (see above), whose loose ends were tightened with elastic bands or sticky tapes. The plates were then kept at 4°C for long-term storage for up to a few months.

### Competent and ultracompetent *E. coli* cells

DH5α competent cells were prepared in house with the protocol detailed below, whereas SURE-2 and TOP10 competent cells were purchased from Agilent (Cat. 200152) and Thermo Fisher Scientific (Cat. C404010), respectively. For comparison, XL10-Gold ultracompetent cells were also purchased from Agilent (Cat. 200314). Transformation of SURE-2, TOP10 and XL10-Gold competent cells were carried out as for DH5α competent cells, but two modifications were made, per manufacturers’ instructions: 1) 10-min incubation on ice with 24 mM β-mercaptoethanol prior to addition of DNA and 2) use of 100 μl NZY^+^ broth (see the recipe below) instead of the SOC medium after the heat-shock step. Compared to DH5α competent cells that had been stored in a liquid nitrogen tank for over 12 years, XL10-Gold ultracompetent cells yielded about twice more colonies. For many plasmids subjected to site-directed mutagenesis, DH5α competent cells prepared in house were adequate. For transformation with SOC, NYZ^+^ and SON media (see the recipes below) did not yield significant difference. Notably, 10-min incubation on ice with 24 mM β-mercaptoethanol prior to addition of DNA did not appear to enhance (but rather decreased) the transformation efficiency with DH5α competent cells prepared below.

DH5α competent cells were prepared with an in-house protocol modified from an early version that we used previously [42,52]. Large-batch preparation of such competent cells is both time-saving and economical, as they can be stored in a liquid nitrogen tank for over 12 years with minimal loss of competency. The protocol described below is also efficient for transformation of many other commonly used bacterial strains, but the impact of long-term storage has not been tested for these strains.

For preparation of DH5α competent cells, a water bath shaker was set up in a cold room and the temperature was adjusted to 18°C. Sterile plastic pipette, pipette tips, 250 ml centrifugation bottles and 2-ml sample vials were cooled down in the cold room. To avoid microbial contamination, the following three steps were carried out close to a benchtop gas flame. DH5α cells were streaked out from a glycerol stock (kept at -80°C) onto a LB agar plate (without any antibiotics) and were grown overnight in a 37°C incubator. With a sterile 200-μl pipette tip, 20-30 colonies (with the diameter of 2∼3 mm) were inoculated into 1 liter of sterile SOB in an autoclaved 2-1 Erlenmeyer flask (Pyrex) covered with 8 layers of cheese cloth and then with one sheet of aluminum foil. After inoculation, the 8-layer cheese cloth and aluminum foil were put back to cover the top of the flask securely. For aeration, the aluminum foil was punched a few times with a clean tip. The flask was shaken in an Innova 3100 water-bath shaker (New Brunswick Scientific) for 40-60 h at 200∼250 rpm until OD_600_ reached 0.4∼0.8 (0.6 would be the best). Afterwards, the flask was cooled down on ice for 10 min; at the same time, the TB solution (see the recipe below) was also pre-chilled on ice.

The following two steps were carried out close to a benchtop gas flame whenever possible. The bacterial culture was transferred into four 250-ml sterile Corning® centrifuge bottles (Sigma, Cat. CLS430776) for centrifugation at 3,000 pm and 4°C for 10 min. The supernatant was decanted and the four cell pellets were suspended in 300 ml of TB by pipetting up and down with a cold and sterile 25 ml plastic pipette; at this step, no vortexing was allowed as it might damage the cells. The cell suspension was kept on ice for 10 min before centrifugation at 3,000 rpm and 4°C for 10 min; it is fundamental to cool down the centrifuge to 4°C prior to centrifugation. Next to a benchtop gas flame, the supernatant was decanted, and the pellet was put back on ice for suspension in 80 ml of a cold TB solution by pipetting up and down with a cold and 10-ml sterile plastic pipette; again, no vortexing was allowed at this step. The cell suspension was kept on ice for 10 min and 6 ml DMSO was slowly added to the cell suspension while the centrifuge bottle was being gently swirled on ice. 2-ml aliquots were pipetted into prechilled 2-ml sterile Sarstedt microtubes (Cat. 72.694.006), which were tightly screw-capped and dropped immediately into a liquid nitrogen bath for flash-freezing. The competent cells were stored in a liquid nitrogen tank for long-term storage (up to 12 years). For convenience and short-term storage, one vial of 2-ml competent cells could be aliquoted further into 0.1-0.2 ml aliquots in 1.5 ml sterile Eppendorf tubes, which were capped and flash-frozen on dry ice for storage at -80°C for up to a few months.

### Preparation of the transformation buffer (TB), SOC and NYZ^+^ media

For preparation of the TB stock solution, it is essential to use ultrapure reagents (including water) and ultraclean glassware. To prepare 1 liter of the TB stock solution, 3.0 g PIPES, 2.2 g CaCl_2_-2H_2_O and 18.6 g KCl were dissolved with 950 ml Nanopure water in a clean 1-liter glass beaker, which was pre-rinsed several times with Nanopure water. The pH value was adjusted with ∼0.9 ml 6 M HCl to 6.7 and 10.9 g MnCl_2_-4H_2_O was then added. After MnCl_2_-4H_2_O was completely dissolved, the solution was adjusted to 1 liter with additional Nanopure water. The solution may appear a bit pinkish, due to the presence of MnCl_2._ For sterilization, the stock solution was filtered through a 0.2 μm-membrane filter unit containing a (such as Corning® bottle-top vacuum filter system) within a cell culture hood.

To prepare the SOB (super optimal broth) medium (per 1 liter), 20 g tryptone, 5 g yeast extract, 0.5 g NaCl (or 1.7 ml 5 M NaCl) and 186 mg KCl (or 1.25 ml of a 2 M stock solution) were weighted directly into a 2-liter Erlenmeyer glass flask. 980 ml of Nanopure water was added, along with 1 ml 1 M NaOH to adjust the final pH value close to 7.0. After being covered with 8 layers of cheese cloth and a sheet of aluminium foil on the top, the flask was sterilized by autoclaving at 15 PSI (1.05 kg/cm^2^) for 20 min on a liquid cycle. After cooling down, 5 ml 1 M MgSO_4_ and 5 ml 1 M MgCl_2_ were added. These solutions were separately prepared in 100-200 ml stocks with Nanopure water in glass bottles and autoclaved.

To prepare SOC (<u>s</u>uper <u>o</u>ptimal broth with <u>c</u>atabolite repression), 250 ml of the SOB medium was prepared as described above, in an autoclavable 500-ml glass bottle with a crew cap. After cooling down to the room temperature, 2.5 ml 1 M glucose (equivalent to 18%) was added, near a gas flame from a Bunsen burner. The glucose stock solution was prepared with autoclaved Nanopure water and sterilized by filtration.

To prepare 250 ml NYZ^+^ broth, 5 g NZ amine A (Sigma, Cat. C0626), 1.25 g yeast extract and 1.25 g NaCl were weighted directly into a 500-ml glass bottle with a crew cap. 235 ml Nanopure water was added, along with ∼0.25 ml 1 M NaOH to adjust pH to 7.5. The bottle was autoclaved at 15 PSI (1.05 kg/cm^2^) for 20 min on a liquid cycle. After colling down to the room temperature, the following sterile solutions were added: 3.1 ml 1 M MgSO_4,_ 3.1 ml 1 M MgCl_2_ and 5 ml 1 M glucose, while working close to a benchtop gas flame.

An SOC-NYZ^+^ hybrid medium, referred to as the SON medium, was also tested for transformation of DH5α competent cells. To prepare this medium (250 ml), 2.5 g tryptone, 2.5 g NZ amine A (Sigma-Millipore, Cat. C0626), 1.25 g yeast extract, 0.5 g NaCl and 46.5 mg KCl (or 0.25 ml 2 M KCl) were weighted directly into a 500-ml glass bottle with a crew cap. 225 ml deionized water was added, along with ∼0.25 ml 1 M NaOH to adjust the final pH value to 7.5. The bottle was autoclaved at 15 psi (1.05 kg/cm^2^) for 20 min on a liquid cycle. After colling down to the room temperature, the following sterile solutions were added: 2 ml 1 M MgSO_4,_ 2 ml 1 M MgCl_2_ and 2.5 ml 1 M glucose, while working close to a benchtop gas flame. Compared to the widely used SOC medium, the SON medium contains NZ amine A and increased amounts of MgSO_4,_ MgCl_2_ and glucose, with slightly higher pH of 7.5 (instead of 7.0) and a reduced amount of tryptone at 10 g/liter, instead of 20 g/liter.

### Estimation of misincorporation rate

DNA sequence alignment tools in the SnapGene software package were used to analyze Sanger DNA sequencing data. Sequence quality was manually assessed from the corresponding sequencing chromatograms, and the aligned sequences were visually inspected to identify unwanted mutations. To calculate the misincorporation rate, the number of unwanted mutations was divided by the total number of the sequenced nucleotides whose corresponding sequencing chromatograms were inspected and determined to be of high-quality.

## Supporting information

Manuscript text and figures

## Acknowledgement

This work was supported by funds from Canadian Institutes of Health Research (CIHR), Natural Sciences and Engineering Research Council of Canada (NSERC) and Compute Canada (to X.J.Y.).

## Data Availability

The authors confirm that the data supporting the findings of this study are available within the article and its supplementary materials. Original data and the plasmids generated from this study are available from the corresponding author, upon reasonable request.

