## Supplementary material for "P3 site-directed mutagenesis: An efficient method based on primer pairs with 3’-overhangs": Manuscript text and figures

**Legends to Supplementary Figures**

**Fig. S1 P3 mutagenesis for engineering single-site vs multi-site mutations**

A. Cartoon illustrating a pair of primers with 3'-overhangs for single-site mutagenesis.

B. Schematic illustrating two partially complementary primers with 3' overhangs (Figs. 1B & S1A) direct synthesis from nicked newly synthesized DNA strands. One such pair of primers (P1F and P1R, denoted with horizontal arrows) is to engineer a mutation, depicted with red asterisks. In stark contrast, completely complementary primers (Fig. 1A) are unable to do so.

C. Cartoon illustrating two pair of primers with 3'-overhangs for dual-site mutagenesis. Due to its small size of the amplified product, PCR between P1F and P2R may dominate amplification and affect the outcome negatively.

D. Cartoon illustrating three pair of primers with 3'-overhangs for three-site mutagenesis. PCR between P1F and P2R (or P2F and P3R) may dominate the amplification and affect the outcome.

**Fig. S2 Pfu-fly polymerase-mediated P3 mutagenesis to produce BRPF3 mutants**

A. Sequence analysis of three plasmids from engineering the R15W mutant of BRPF3. Pfu-fly was used for P3 mutagenesis. All three plasmids contain large deletions (~ 3 kb). Moreover, plasmids from colonies 1 and 2 also possess small insertions (10-20 bp) at the primer site (data not shown). C1, C2 and C3 denote plasmids from colonies 1, 2 and 3. We sequenced additional colonies and all of them were either wild-type or contained large deletions as C1 and C2.

B. Sanger sequence analysis of three plasmids from generating the R51H mutation of BRPF3. Pfu-fly was used for P3 mutagenesis. All three plasmids carry the expected mutations, so the efficiency was 100%. The striking variability of the method in engineering this and R15W (see panel A) indicates that further improvement is required to extend the method for certain special cases.

**Fig. S3 Efficient construction of JADE mutants through P3 site-directed mutagenesis.**

A. Sanger sequence analysis of 7 plasmids from engineering the I76A and L77A mutants of JADE2 with only one pair of primers. Five are for the I76A substitution, one encoded L77A substitution and one was a mixed clone of both mutants, so the mutagenesis efficiency was 100%.

B. Sanger sequence analysis of three plasmids from engineering the E463* truncation mutant of JADE3. All three carried the expected mutation, so the mutagenesis efficiency was 100%.

**Fig. S4 P3 site-specific mutagenesis to generate EPC1 and EPC2 mutants**

A. Domain organization of EPC1 and EPC2. Like EPC2, ECP1 is composed of to an N-terminal part (N) for interacting with KAT7 [32] and two EPC-like motifs (I and II) [32]. Five nonsense variants are illustrated, with three for EPC1 (Q355*, W425* and E543*) and two for EPC2 (E468* and E542*), where asterisks denote stop codons.

B. Sanger sequence analysis of 9 plasmids from engineering the Q355* variant of EPC1. Four were correct, so the efficiency was 44.4%. The same mutagenesis reaction mixture was used for transformation of DH5α and SURE2. In terms of mutagenesis efficiency, DH5α appeared to be better than SURE2, which is lack of some genes needed for DNA recombination and repair.

C. Sanger sequence analysis of 9 plasmids from engineering the W425* variant of EPC1. Three could not be sequenced and three were correct, so the efficiency was 33%.

D. Sanger sequence analysis of 4 plasmids from engineering the E543* variant of EPC1. All four were correct, so the efficiency was 100%.

E. Sequence chromatograms of 3 plasmids from engineering the E468* variant of EPC2. Two were correct and the third was a mixed clone, so the efficiency was 2.5/3=83%.

F. Sanger sequence analysis of 16 plasmids from engineering the E542* variant of EPC2. Among them, 14 contained the mutation, with one harboring an unwanted T>A substitution 49 downstream from the designed mutation site and another was a mixed clone, so the efficiency was 13.5/16=84%.

**Fig. S5 P3 site-directed mutagenesis to engineer HDAC4 and HDAC5 mutants.**

A. Domain organization of HDAC4 and its paralog HDAC5. HDAC4 is composed of to an N-terminal motif for MEF2 binding, three sites for 14-3-3 binding shown as small rectangles marked with the dark letter S, an NLS (nuclear localization signa), a serine-rich motif shown as a small rectangle marked with red letter S, a deacetylase domain and an NES (nuclear export sequence) [53]. Four missense variants are illustrated, with two for HDAC4 (P248L and P248A) and two for HDAC5 (S318A and S322A). The former two were identified in patients with a developmental disorder [49], whereas the latter two are artificial mutants to investigate function of the serine-rich motif [53].

B. Sanger sequence analysis of 6 plasmids from engineering the P248L and P248A variants of GFP-HDAC4. Note that only one primer pair was used for the mutagenesis reaction, with the strategy listed in Fig. 1D. Among 10 candidate plasmids that were sequenced, five contained P248L but one was for P248A, so the mutagenesis efficiency was 60%. For FLAG-tagged HDAC4, four were sequenced, with three being P248L and one for P248A, so the mutagenesis efficiency was 100%.

C. Sanger sequence analysis of 4 plasmids from generating the S318S variant of HDAC5. One was correct, so the efficiency was 25%.

D. Sanger sequence analysis of 4 plasmids from engineering the S322A variant of HDAC5. Two contained the mutation but one was a mixed clone, so the efficiency was 1.5/4=~40%.

**Fig. S6 Pfu-fly based P3 mutagenesis for the spike protein of SARS-COV-2.**

A. Sequence chromatograms of four plasmids sequenced for engineering the L455S/F456L and F456L mutants of the spike protein of the Omicron variant. Pfu-fly polymerase and only one pair of primers was used for engineering the L455S/F456L and F456L mutants of the Omicron spike protein. Among the four plasmids sequenced, three carried the L455S/F456L substitutions and the fourth encoded F456L, so the mutagenesis efficiency was 100%.

B. Sanger sequence analysis of plasmids from engineering the V1104 mutant of the spike protein of the D614G variant. Pfu-fly polymerase was used for P3 mutagenesis. Among four plasmids sequenced, three carried the expected substitution and the fourth showed poor sequence quality (not shown here), so the mutagenesis efficiency was 75%.

**Fig. S1**

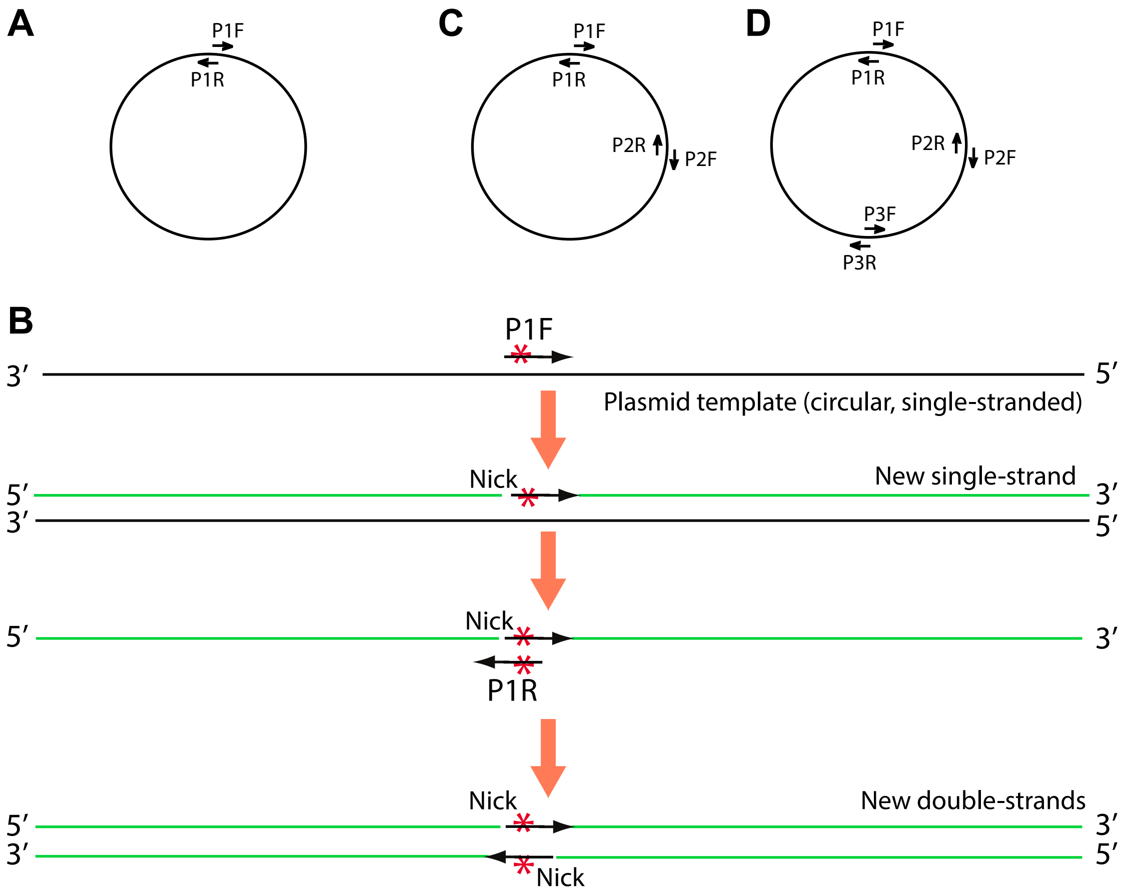
A
P1F
P1R
C
P1F
P1R
P2R
P2F
D
P1F
P1R
P2R
P2F
P3F
P3R
B
P1F
3′
5′
Plasmid template (circular, single-stranded)
Nick
New single-strand
5′
3′
3′
5′
Nick
5′
3′
P1R
Nick
New double-strands
5′
3′
3′
5′
Nick

**Fig. S2**

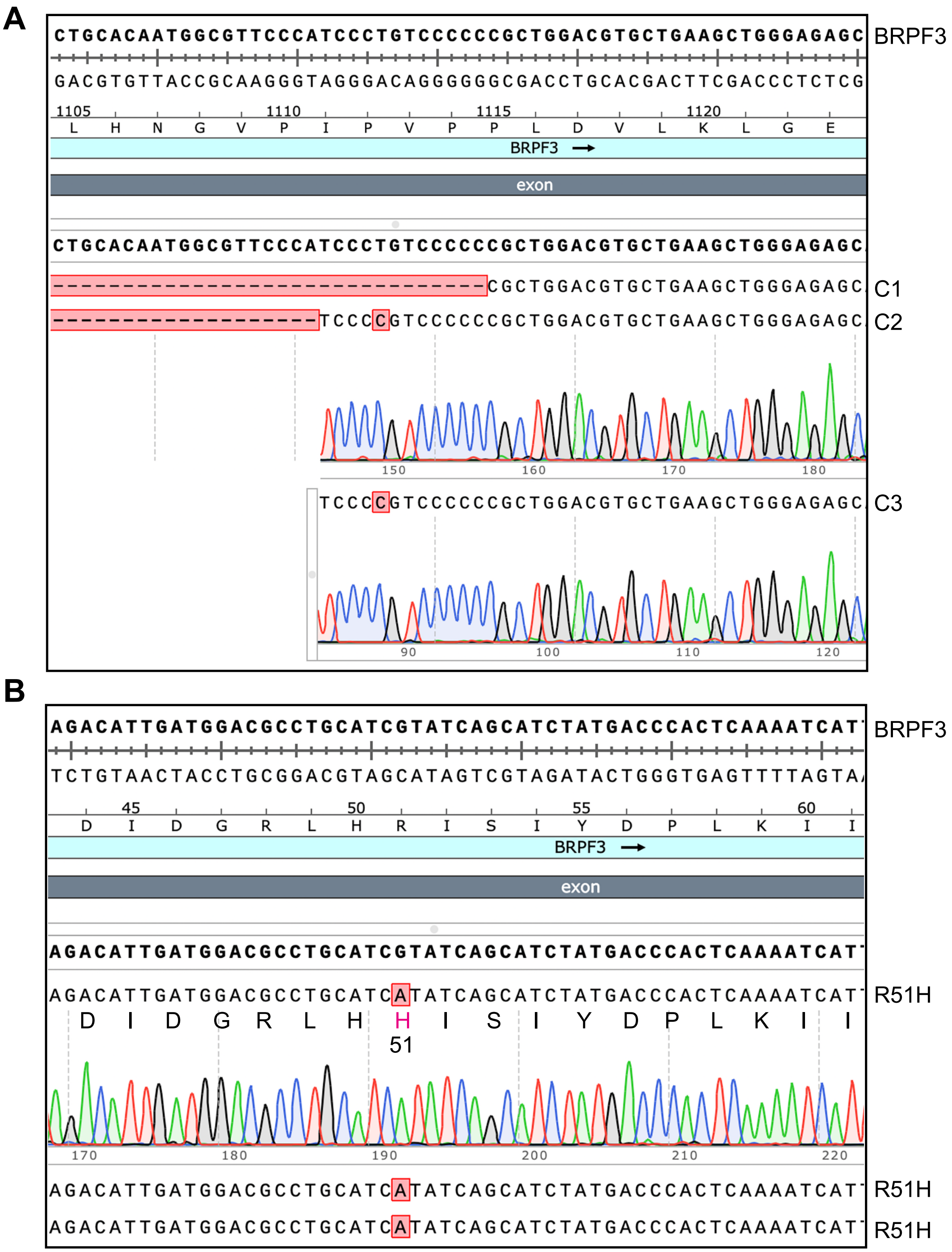
A
CTGCACAATGGCGTTCCCATCCCTGTCCCCCCGCTGGACGTGCTGAAGCTGGGAGAGC
BRPF3
GACGTGTTACCGCAAGGGTAGGGACAGGGGGGCGACCTGCACGACTTCGACCCTCTCG
1105 1110 1115 1120
L H N G V P I P V P P L D V L K L G E
BRPF3
exon
CTGCACAATGGCGTTCCCATCCCTGTCCCCCCGCTGGACGTGCTGAAGCTGGGAGAGC
CGCTGGACGTGCTGAAGCTGGGAGAGC
C1
TCCCCGTCCCCCCGCTGGACGTGCTGAAGCTGGGAGAGC
C2
150 160 170 180
TCCCCGTCCCCCCGCTGGACGTGCTGAAGCTGGGAGAGC
C3
90 100 110 120
B
AGACATTGATGGACGCCTGCATCGTATCAGCATCTATGACCCACTCAAAATCAT
BRPF3
TCTGTAACTACCTGCGGACGTAGCATAGTCGTAGATACTGGGTGAGTTTTAGTA
45 50 55 60
D I D G R L H R I S I Y D P L K I I
BRPF3
exon
AGACATTGATGGACGCCTGCATCGTATCAGCATCTATGACCCACTCAAAATCAT
AGACATTGATGGACGCCTGCATCATATCAGCATCTATGACCCACTCAAAATCAT
R51H
D I D G R L H H I S I Y D P L K I I
51
170 180 190 200 210 220
AGACATTGATGGACGCCTGCATCATATCAGCATCTATGACCCACTCAAAATCAT
R51H
AGACATTGATGGACGCCTGCATCATATCAGCATCTATGACCCACTCAAAATCAT
R51H

### Fig. S3

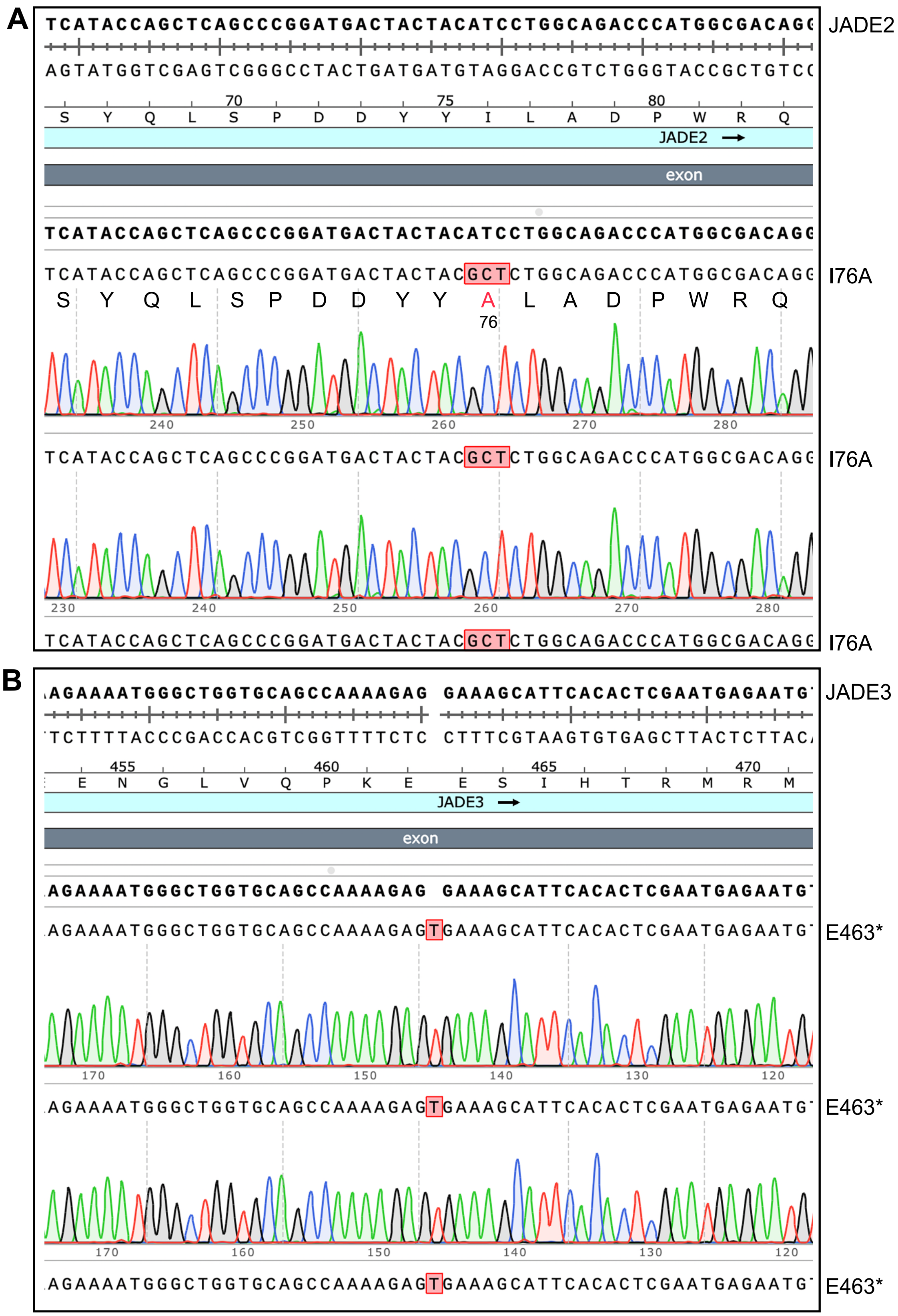
A
TCATACCAGCTCAGCCCGGATGACTACTACATCCTGGCAGACCCATGGCGACAGG
JADE2
AGTATGGTCGAGTCGGGCCTACTGATGATGTAGGACCGTCTGGGTACCGCTGTCC
70
75
80
S Y Q L S P D D Y Y I L A D P W R Q
JADE2
exon
TCATACCAGCTCAGCCCGGATGACTACTACGCTCTGGCAGACCCATGGCGACAGG
I76A
S Y Q L S P D D Y Y A L A D P W R Q
76
I76A
I76A
B
AGAAAATGGGCTGGTGCAGCCAAAAGAG GAAAGCATTCACACTCGAATGAGAATG
JADE3
TCTTTTACCCGACCACGTCGGTTTTCTC CTTTCGTAAGTGTGAGCTTACTCTTAC
455
460
465
470
E N G L V Q P K E E S I H T R M R M
JADE3
exon
AGAAAATGGGCTGGTGCAGCCAAAAGAGTGAAAGCATTCACACTCGAATGAGAATG
E463*
E463*
E463*

**Fig. S4**

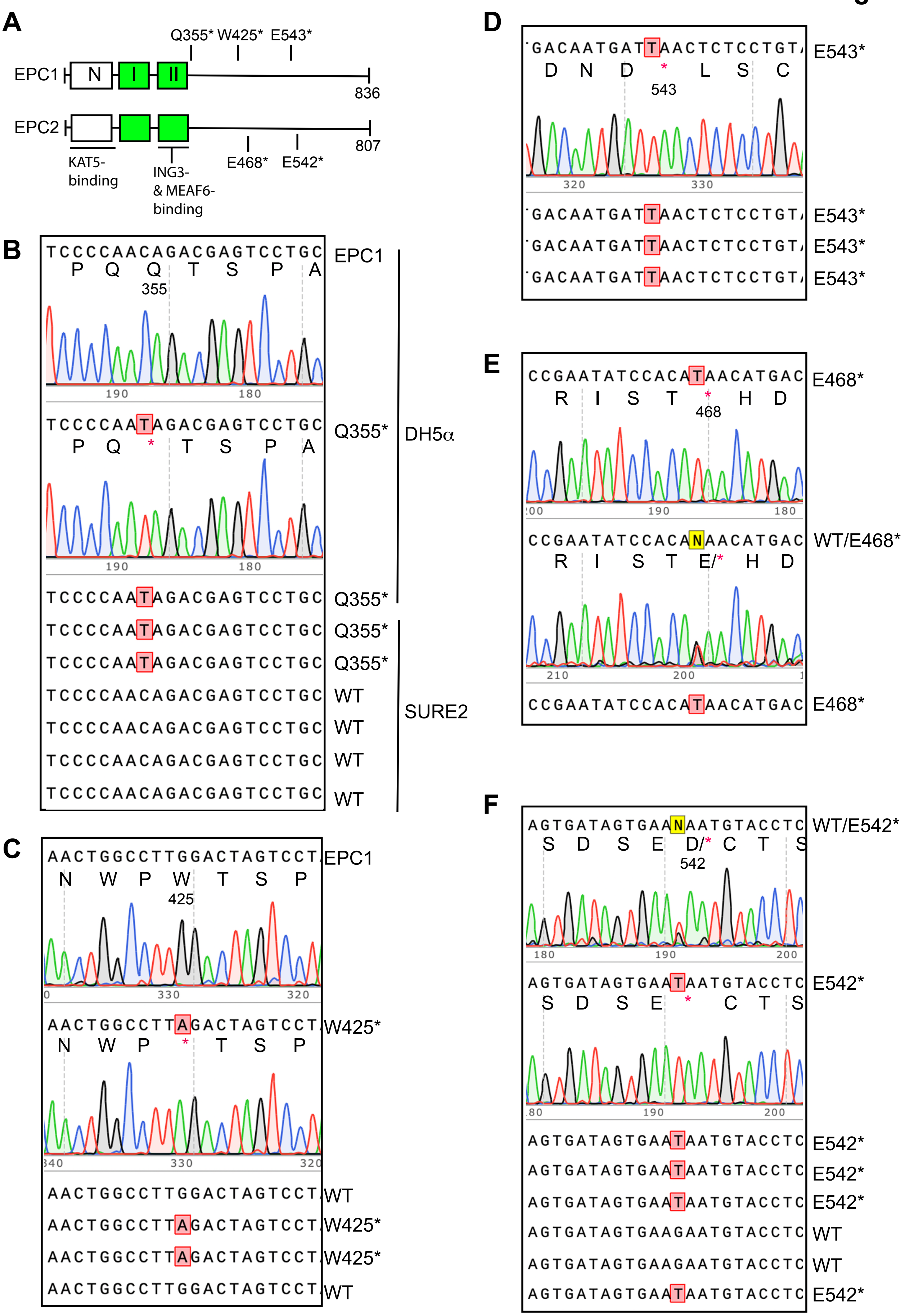
A
Q355* W425* E543*
EPC1
N
I
II
836
EPC2
807
KAT5-
binding
ING3-
& MEAF6-
binding
E468* E542*
B
TCCCCAACAGACGAGTCCTGC
P Q Q T S P A
355
EPC1
TCCCCAATAGACGAGTCCTGC
P Q * T S P A
Q355*
DH5α
TCCCCAATAGACGAGTCCTGC Q355*
TCCCCAATAGACGAGTCCTGC Q355*
TCCCCAATAGACGAGTCCTGC Q355*
TCCCCAACAGACGAGTCCTGC WT
TCCCCAACAGACGAGTCCTGC WT
TCCCCAACAGACGAGTCCTGC WT
TCCCCAACAGACGAGTCCTGC WT
SURE2
C
AACTGGCCTTGGACTAGTCCT
N W P W T S P
425
EPC1
AACTGGCCTTAGACTAGTCCT
N W P * T S P
W425*
AACTGGCCTTGGACTAGTCCT WT
AACTGGCCTTAGACTAGTCCT W425*
AACTGGCCTTAGACTAGTCCT W425*
AACTGGCCTTGGACTAGTCCT WT
D
GACAATGATTAACTCTCCTGT
D N D * L S C
543
E543*
GACAATGATTAACTCTCCTGT E543*
GACAATGATTAACTCTCCTGT E543*
GACAATGATTAACTCTCCTGT E543*
E
CCGAATATCCACATAACATGAC
R I S T * H D
468
E468*
CCGAATATCCACANAACATGAC
R I S T E/* H D
WT/E468*
CCGAATATCCACATAACATGAC E468*
F
AGTGATAGTGAANAATGTACCTC
S D S E D/* C T S
542
WT/E542*
AGTGATAGTGAATAATGTACCTC
S D S E * C T S
E542*
AGTGATAGTGAATAATGTACCTC E542*
AGTGATAGTGAATAATGTACCTC E542*
AGTGATAGTGAATAATGTACCTC E542*
AGTGATAGTGAAGAATGTACCTC WT
AGTGATAGTGAAGAATGTACCTC WT
AGTGATAGTGAATAATGTACCTC E542*

**Fig. S5**

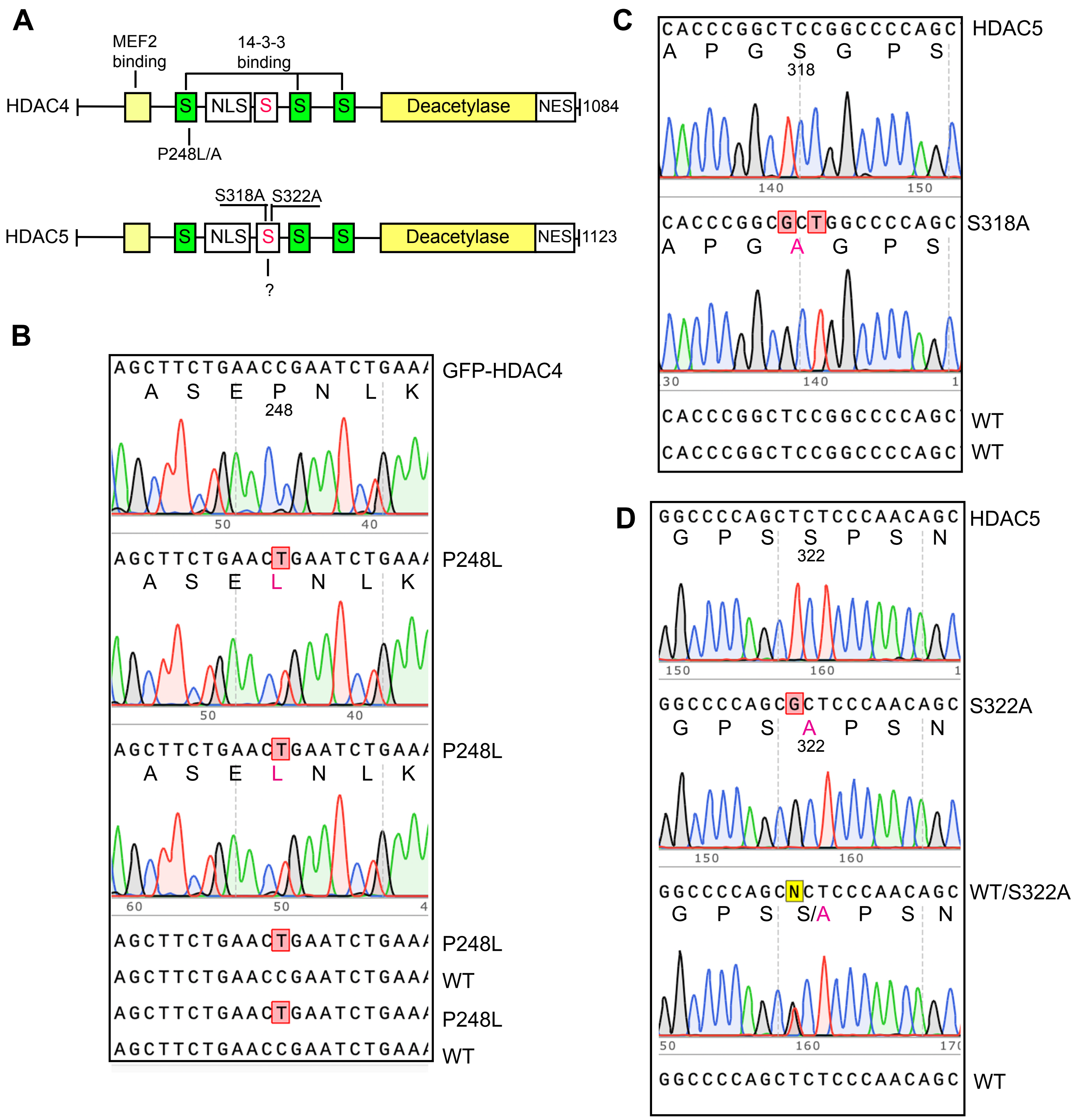
A
MEF2 binding
14-3-3 binding
HDAC4
S
NLS
S
S
S
Deacetylase
NES
1084
P248L/A
S318A
S322A
HDAC5
1123
?
B
AGCTTCTGAACCGAATCTGAA
A S E P N L K
248
GFP-HDAC4
P248L
A S E L N L K
WT
C
CACCCGGCTCCGGCCCCAGC
A P G S G P S
318
HDAC5
S318A
A P G A G P S
D
GGCCCCAGCTCTCCCAACAGC
G P S S P S N
322
S322A
G P S A P S N
WT/S322A
G P S S/A P S N

### Fig. S6

A

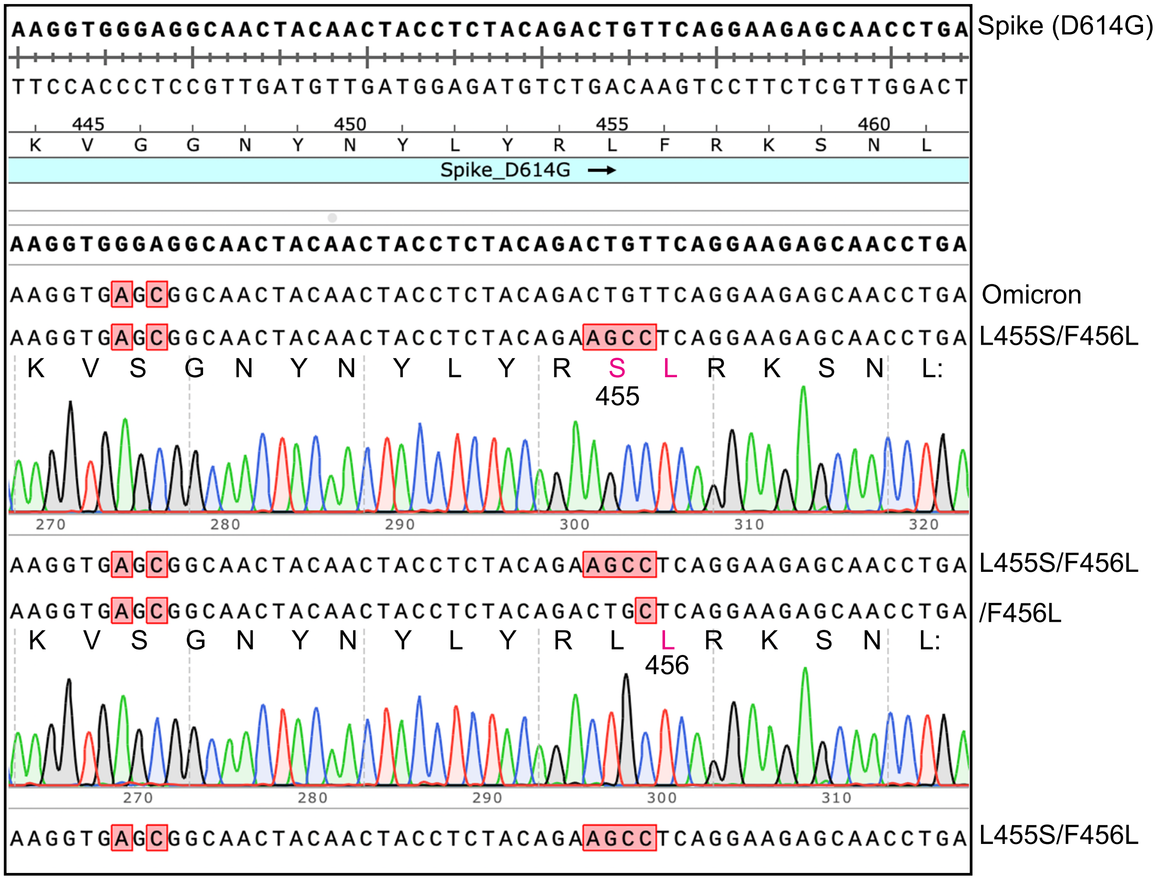

Spike (D614G)
Spike_D614G
Omicron
L455S/F456L
455
L455S/F456L
/F456L
456
L455S/F456L


B

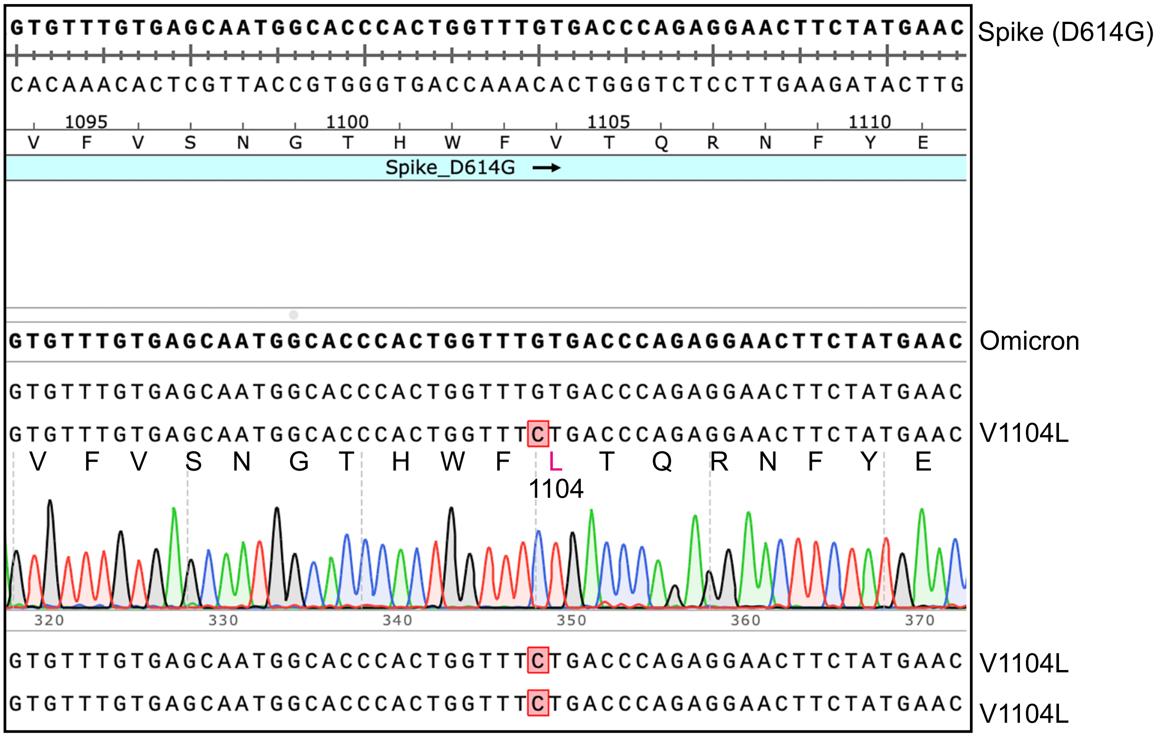

Spike (D614G)
Spike_D614G
Omicron
V1104L
1104
V1104L
V1104L
